# H9N2 avian influenza virus in Bangladesh: three genetically distinct co-circulating subgroups retain a largely conserved antigenic phenotype

**DOI:** 10.64898/2026.11.13.766375

**Authors:** Rebecca Daines, Jean-Remy Sadeyen, Mahmudul Hasan, Tridip Das, Rashed Mahmud Sizer, Md Helal Uddin, Paritosh Kumar Biswas, Md Abdus Samad, Md Ahasanul Hoque, Anne Conan, Joshua Lynton-Jenkins, Ashley Banyard, Ian Brown, Tom Lewis, Jayna Raghwani, Guillaume Fournie, Damer Blake, Fiona Tomley, Munir Iqbal

## Abstract

Low pathogenic avian influenza (LPAI) H9N2 viruses are endemic in poultry across Bangladesh, causing substantial economic losses and posing a persistent zoonotic risk. Although genetic surveillance has documented ongoing viral evolution, the relationship between genetic diversification and antigenic change remains poorly understood. This study investigated the genetic diversity, reassortment, and antigenic characteristics of contemporary H9N2 viruses circulating in Bangladesh.

Phylogenetic analysis of 219 sampled sequences revealed three genetically distinct monophyletic haemagglutinin (HA) G-lineage sub-groups, each characterised by unique amino acid substitutions. In contrast, neuraminidase (NA) displayed similar genetic diversity without distinct phylogenetic clusters. A dual-metric analysis of paired HA and NA phylogenies identified 20 putative reassortant H9N2 viruses.

Further analysis showed that genetic diversity was geographically structured; however, all three HA subgroups co-circulated across multiple regions, and some live bird markets contained representatives of all three groups simultaneously, suggesting that live bird markets may facilitate mixing between genetically distinct viruses, and create opportunities for reassortment among otherwise geographically separated strains. Despite substantial genetic diversity and reassortment, all viruses remained antigenically similar based on haemagglutination inhibition and virus neutralisation assays. However, a representative strain from one subgroup induced significantly lower homologous and heterologous antibody titres, suggesting reduced immunogenicity.

Our findings demonstrate a notable disconnect between genetic evolution and antigenic change in Bangladeshi H9N2 viruses. While reassortment and genetic diversification continue to shape viral populations, antigenic properties remain largely conserved, supporting the continued relevance of current vaccine strategies while highlighting the importance of ongoing genetic and antigenic surveillance.

**Importance:** Low pathogenic avian influenza (LPAI) H9N2 viruses remain endemic in Bangladesh and continue to evolve through genetic diversification and reassortment. This study demonstrates that, despite the emergence of three distinct HA genetic subgroups and frequent reassortment events, contemporary Bangladeshi H9N2 viruses remain antigenically conserved. These findings reveal an important disconnect between genetic evolution and antigenic change, indicating that genetic divergence alone may not predict vaccine effectiveness. The co-circulation of multiple viral subgroups within live bird markets highlights their role as hubs for viral mixing and reassortment, emphasizing the need for targeted surveillance in these settings. While current vaccine strategies are likely to remain effective against circulating strains, the identification of a subgroup with reduced immunogenicity underscores the importance of continued monitoring of both genetic and phenotypic traits. Together, these findings provide critical evidence to support risk assessment, vaccine evaluation, and avian influenza control programmes in Bangladesh and beyond.

## Introduction

H9N2 influenza is a low pathogenic avian influenza virus (AIV) that has become endemic in poultry populations across large parts of Asia, the Middle East, and increasingly other regions [1]. Although it is generally associated with low mortality in birds, its persistent circulation in commercial and backyard poultry systems generates substantial and sustained economic losses by its effects on poultry performance rather than mass mortality events; infected flocks typically exhibit reduced egg production, slower growth rates, impaired feed conversion efficiency, and increased vulnerability to secondary infections, all of which directly reduce farm profitability [2, 3].

Bangladeshi poultry farms adopt vaccination strategies to control AIV spread and disease through a combination of imported commercial inactivated vaccines and locally produced autogenous/local inactivated vaccines [4]. Because the Bangladesh Department of Livestock Services (DLS) approves specific commercial products for distribution, a few key seed strains dominate field usage [5]. However, field studies and challenge trials in Bangladesh demonstrate that current H9N2 vaccines protect against clinical disease and mortality, but do not provide sterilising immunity or completely prevent shedding [5–7]. Absent or inefficient protection, coupled with rapid mutation rates, promotes the emergence of distinct antigenic variants able to escape vaccine-induced and pre-existing immunity [8–10]. As such, H9N2 continues to circulate despite long-term vaccination programs.

H9N2 has over time diversified into two major geographical lineages, A (North American) and B (Eurasian), as classified by the WOAH/FAO OFFLU lineage classification system [11]. Lineage A is restricted mainly to wild waterfowl and shorebird reservoirs with very occasional spillover to domestic turkeys or small poultry holdings. Unlike Lineage B, Lineage A has not established sustained, high-density commercial poultry endemicity, resulting in less antigenic drift compared to Eurasian poultry strains [11, 12]. The B lineage is further divided into two sub-lineages due to extensive diversity, sub-lineage B (previously Y280/BJ94), dominating East and

Southeast Asia, and sub-lineage G (G1-derived) dominating Central and South Asia, the Middle East, and North and East Africa. Driven by regional trade, intense agricultural density, and vaccination selection pressure, these clades dominate East Asian poultry and regularly seed reassortant viruses into wild and domestic birds [11, 13]. Similarly, the G lineages underwent spatial compartmentalisation across the Middle East and the Indian subcontinent. Local movement of poultry and continuous trade routes have sustained unique regional sub-clades across countries like India, Pakistan, Iran, and expanding into parts of Africa [14].

The genetic profile of prevailing H9N2 strains in Bangladesh is largely attributed to long-term longitudinal studies utilising sporadic outbreak reports or small-scale sampling programs. However, a larger genetic profile with respective antigenic phenotyping within a single sampling period remains absent. Therefore, this study sought to genetically characterise prevailing strains of H9N2 across Bangladesh and investigate antigenic divergence between them to contribute to improved future vaccine success.

## Results

### Three genetically diverse monophyletic subgroups of the H9N2 G-lineage haemagglutinin (HA) co-circulate across Bangladesh

Haemagglutinin (HA) and neuraminidase (NA) nucleotide sequences acquired from whole genome isolation across Bangladesh (accession numbers Table S1) were phylogenetically analysed for inherent genetic diversity of H9N2 across Bangladesh (Figure 1 & Figure 2).

**Figure 1:**
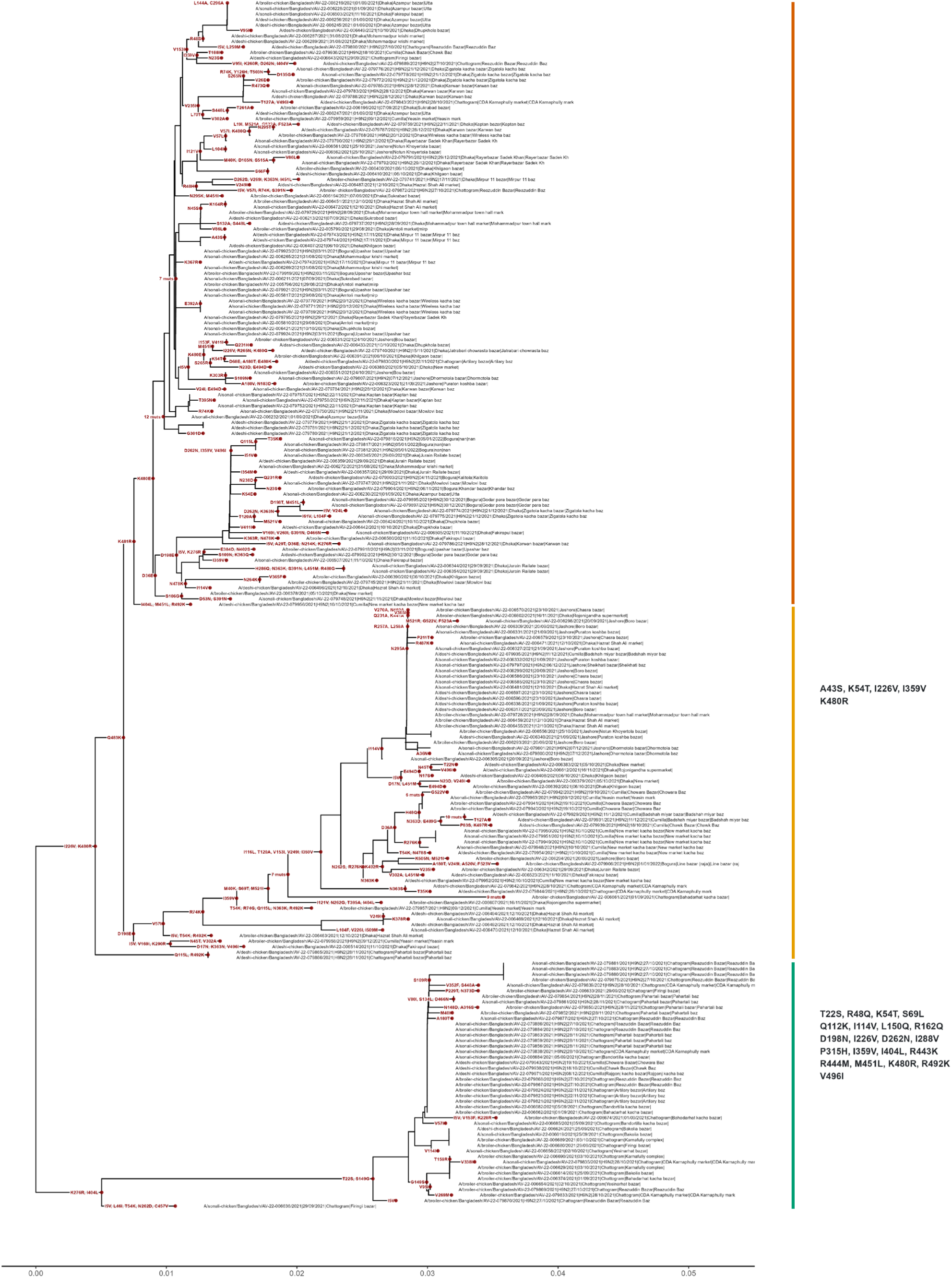
Neighbour-Joining phylogenetic tree of trimmed mature haemagglutinin (HA) amino acid sequences (n = 219) of isolates collected across Bangladesh, with synapomorphic amino acid mutations indicated at the nodes. Scale indicates rate of genetic distance with 0.01 corresponding to 1% difference in amino acid sequence. Tree generated in RStudio [15] with ape [16], phangorn [17], ggtree [18] & tidyverse [19] packages. Each monophyletic group is independently coloured (top = orange, middle = yellow, bottom = green). Conserved mutations (≥95%) from the most ancestral node are listed per branch.

**Figure 2:**
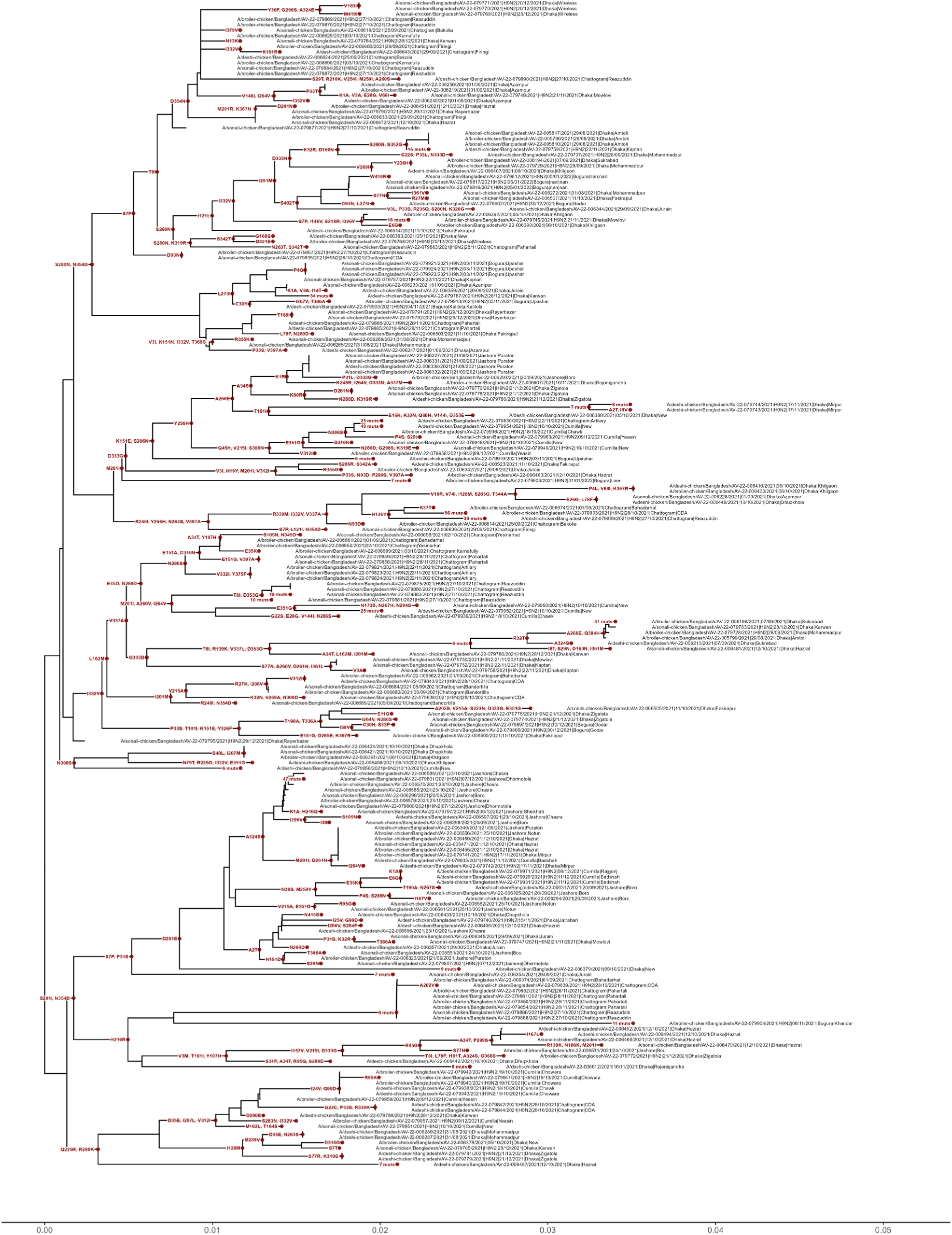
Neighbour-Joining phylogenetic tree of trimmed mature neuraminidase (NA) amino acid sequences (n = 215) of isolates collected across Bangladesh, with synapomorphic amino acid mutations indicated at the nodes. Scale indicates rate of genetic distance with 0.01 corresponding to 1% difference in amino acid sequence. Tree generated in RStudio [15] with ape [16], phangorn [17], ggtree [18] & tidyverse [19] packages.

The HA sequences distinctly clustered into three monophyletic branches of the G-lineage, characterised by a significant amount of synapomorphic amino acid mutations conserved within the branch. For downstream analysis, each monophyletic branch was coloured (top = orange, middle = yellow, bottom = green) (Figure 1). The uppermost and middle monophyletic branches contain isolates from all sampled regions, Bogura, Chattogram, Cumilla, Dhaka and Jashore. However, the lowermost branch only contains isolates from Chattogram and Cumilla. This suggests synergistic harmony without competition between the divergent strains, some degree of independent evolution through ecological or geographical segregation or similar transmission and fitness advantages despite genetic distinction.

The defining mutations differ considerably among the green, yellow and orange monophyletic groups; mutations conserved in ≥95% of strains within each group are indicated (Figure 1). The green monophyletic group contained 21 conserved mutations, including 17 unique substitutions and four shared with the yellow monophyletic group. The yellow monophyletic group contained one additional unique amino acid substitution. Due to the high variability of the orange monophyletic group, no unique mutations were conserved in ≥95% of strains.

The phylogenetic arrangement of NA sequences portrays little to no monophyletic homology compared to HA with multiple less dynamic smaller branch clusters (Figure 2). Where HA presented some unique geographic isolation of the basal branch, all regions are widespread throughout the tree. This could suggest varied degrees of evolution or reassortment of NA compared to HA.

### Haemagglutinin (HA) and neuraminidase (NA) of Bangladeshi H9N2 viruses undergo varying rates of genetic evolution and potential reassortment

To investigate comparative phylogenetic alignment of each gene, a tanglegram of HA and NA sequences was generated (Figure 3). Visual inspection of the fixed topology tanglegram revealed profound phylogenetic incongruence and structural uncoupling between the HA and NA genes, with lines exhibiting heavy crisscrossing across the trees. This pattern strongly indicates frequent genetic reassortment between co-circulating subgroups, demonstrating independent evolutionary trajectories of HA and NA. There is significant group dispersion within the yellow HA monophyletic group, with the highest degree of visual scattering of line connections. The widespread fanning of lines without forming thicker dense bundle lines indicates widespread topological incongruence. In contrast, the green monophyletic HA group features some prominent bundled lines to the upper and middle of the NA tree. These parallel clusters show a significant proportion of the strains share highly consistent positions across both trees. Nonetheless, some individual lines do diverge throughout the NA tree which may more likely suggest reassortment. Finally, the orange monophyletic HA group displays a combination of both trends with some thicker line bundles towards the bottom of the NA tree, but the vast majority of connections are widely distributed. Each monophyletic group of HA displays varied degrees of topological congruence to its NA counterparts which may be attributed by their geographical isolation and therefore independent evolution.

**Figure 3:**
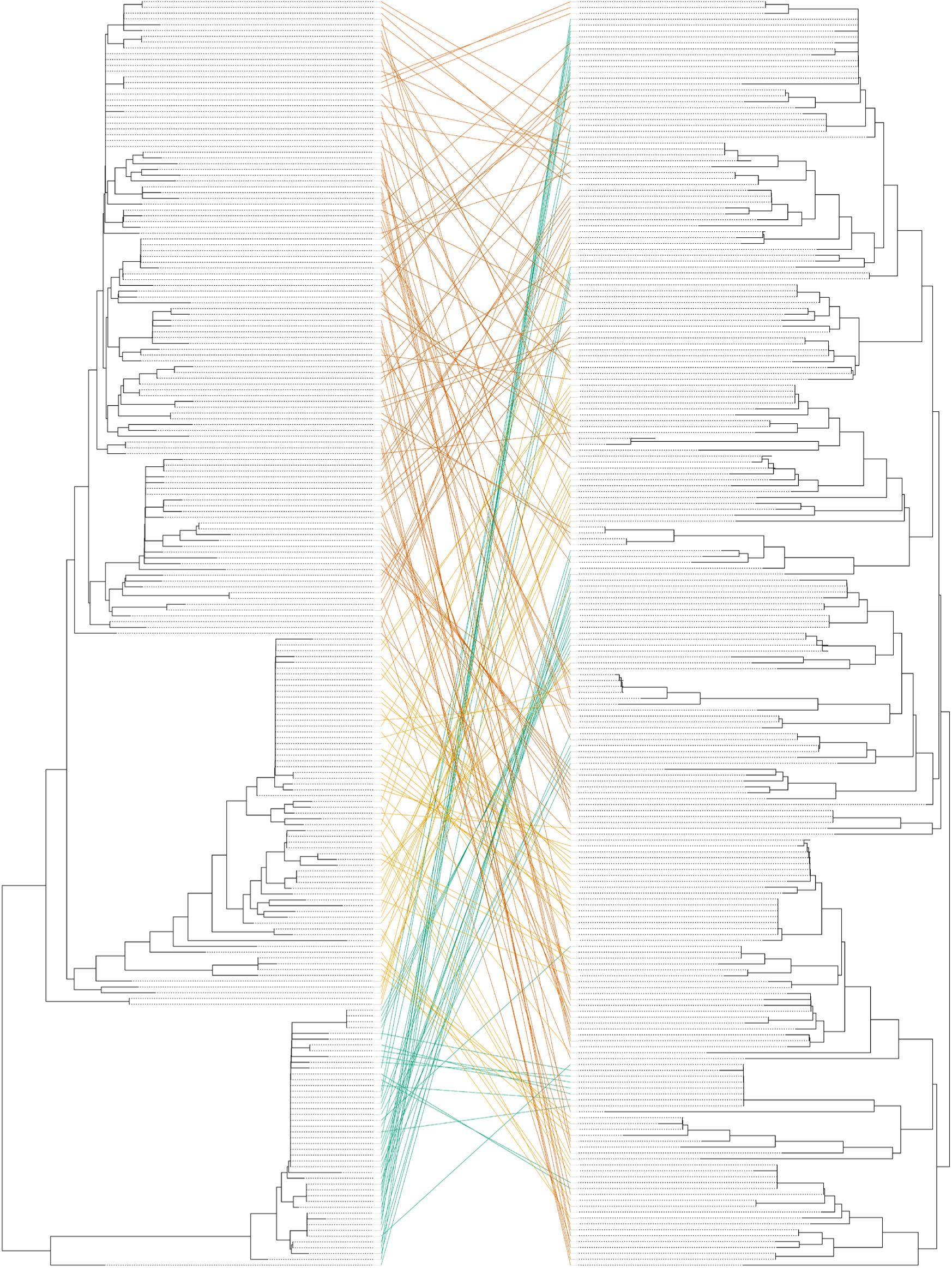
Tanglegram illustrating the topological congruence and segment reassortment between the Neighbour-Joining phylogenetic trees of haemagglutinin (HA) (left) and neuraminidase (NA) (right) of Bangladeshi poultry H9N2 isolates. The lines connecting the HAs and NAs of the same strain are coloured according to the monophyletic groups of HA (Figure 1). Both trees are depicted as cladograms with right-ladderised topology. Tree generated in RStudio [15] with ape [16] and phytools [20] packages.

To further investigate the likelihood of reassortment and quantify segment co-evolution, we calculated a dual-metric threshold system combining composite genetic mismatch scores with k-nearest neighbour (k = 10) topological overlaps (Table S2). Strains were classified into true reassortant (mismatch score > mean + 1.5 standard deviation (StD) and neighbour overlap ≤ 30%), co-evolved/variant neighbour overlap ≥ 60%), or intermediate/drift (neighbour overlap 31-59%). Of 215 strains, n=20 are predicted true reassortants, n=2 co-evolved/variant pairings and n=193 intermediate or drifted gene segments. Of the true reassortants, n=17 of the HAs are located in the lowermost monophyletic group (green), n=2 in the middle group (yellow) and n=1 in the uppermost group. Of the co-evolved/variants, n=1 is located in the lowermost monophyletic group and n=1 in the middle group.

### Monophyletic groups of haemagglutinin (HA) suggest geographically independent evolution with synchronous isolation of each from the same sample site

To further investigate the epidemiological distribution of the genetic diversity of isolates, each monophyletic branch colour (top = orange, middle = yellow, bottom = green) and each strain was plotted according to coordinates of isolation (Figure 4). A total of n=39 sites were sampled across Bangladesh. The top and middle branches with site numbers 1 to 9 correspond to Chittagong, 10 to 14 Jashore, 15 to 17 Cumilla, 18 to 37 Dhaka and 38 to 39 Bogura market clusters (n=5). Of the sites sampled, n=19 sites found isolates from a single monophyletic group, n=18 sites found isolates from two monophyletic groups and n=2 sites found isolates from all three. The green monophyletic group was unique to sites 1, 5, 6 and 9, located in Chittagong, correlate to the phylogenetic analysis of primarily Chittagong and Cumilla sequences. The dominant red monophyletic group naturally coincides with the majority of unique sites, 19, 20, 21, 22, 25, 26, 28, 29, 30, 31, 33, 35, 36 and 37 within Dhaka; site 39, also unique to the dominant red monophyletic group, was further north in Bogura.

**Figure 4:**
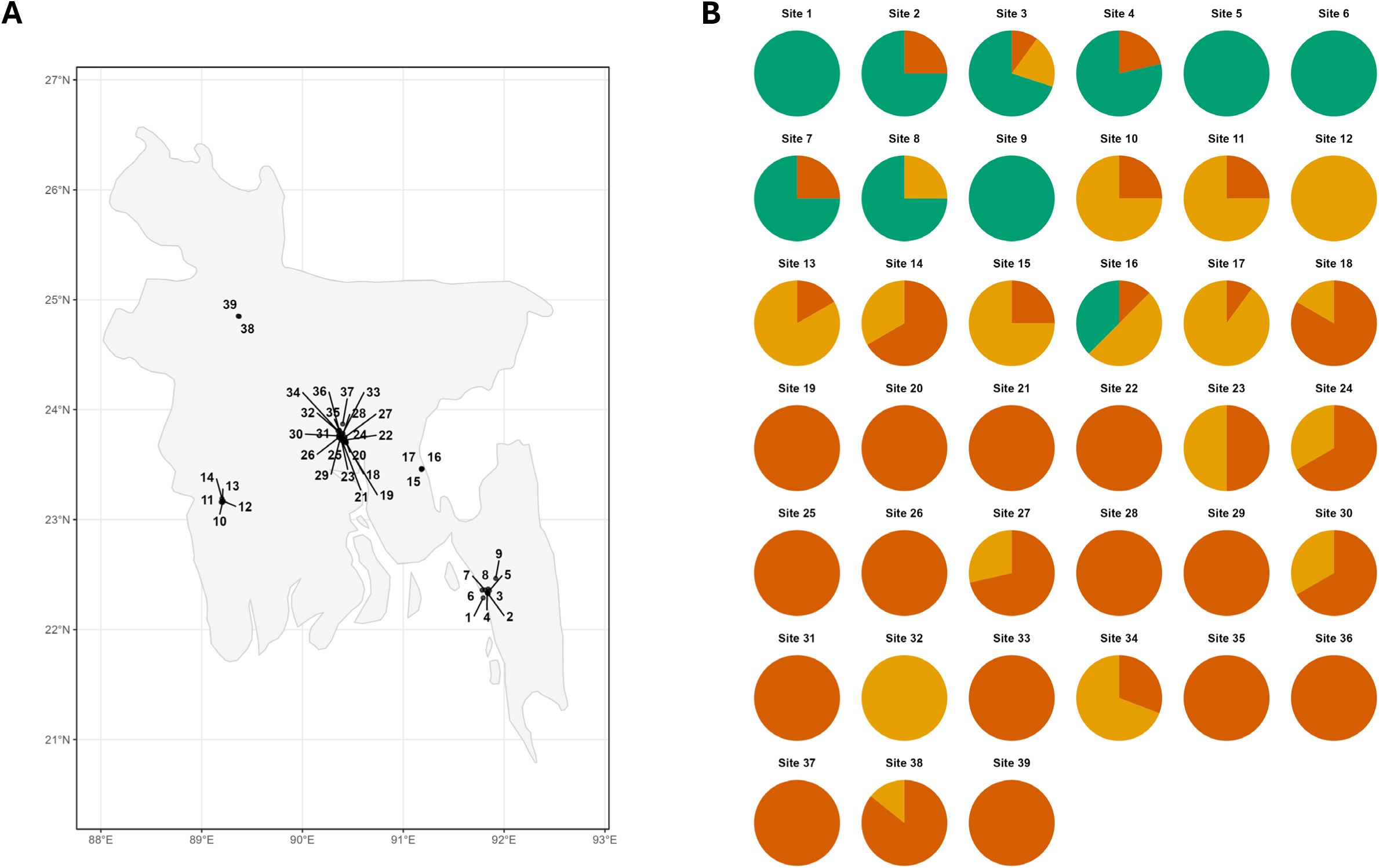
Epidemiological mapping of isolates by numbered site across Bangladesh (**A**) and prevalence of diversity coloured by haemagglutinin (HA) monophyletic subgroup of each numbered site (**B**). Site numbers 1 to 9 correspond to Chittagong, 10 to 14 Jashore, 15 to 17 Cumilla, 18 to 37 Dhaka and 38 to 39 Bogura. Combined image generated in RStudio [15] (ape [16], phangorn [17], tidyverse [19], sf [21], rnaturalearth [22], ggrepel [23], patchwork [24]) packages. Frequency of isolates and coordinates of sites are provided in Table S3.

### Antigenic phenotype is retained across all representative viruses but one presents weak immunogenicity

To explore the antigenic diversity between the HA monophyletic groups, three representative strains, one per group, were selected based upon similarity to the genetic consensus of each group: A/sonali-chicken/Bangladesh/AV-22-079924/2021 – top group (orange) (BNG924); A/sonali-chicken/Bangladesh/AV-22-079948/2021–middle group (yellow) (BNG948); A/deshi-chicken/Bangladesh/AV-22-079943/2021 – bottom group (green) (BNG943). To confirm the strains were both representative of the sequence panel and published data, publicly available H9N2 HA sequences from 2020-2026 were acquired from GISAID (https://gisaid.org/) and phylogenetically aligned with the representative strains (Figure S1). Published sequence data recapitulated the three monophyletic groups and each representative strain aligned into each representative group, suggesting Bangladesh has maintained the distinct co-evolution of HA.

Additionally, each representative was initially genetically analysed for key markers of antigenic divergence of HA and NA; each sequence was comparatively aligned to identify amino acid differences between the strains and percentage variance (Table 1) and prediction of N-linked glycans (Table S4).

**Table 1:** Sequence variant alignment, visualised in MEGAX [26], and coloured by their respective haemagglutinin (HA) monophyletic subgroup where the substitution is unique (**A**) and number of amino acid substitutions with percentage difference (**B**) of the HA G-lineage subgroup candidate viruses and their coupled neuraminidase (NA). Mature H9 HA and H9 N2 NA numbering used.

| Strain | Residue Position (HA) |  |  |  |  |  |  |  |  |  |  |  |  |  |  |  |  |  |  |  |  |  |  |  |  |  |  |
|---|---|---|---|---|---|---|---|---|---|---|---|---|---|---|---|---|---|---|---|---|---|---|---|---|---|---|---|
|  | 22 | 43 | 48 | 54 | 69 | 112 | 114 | 115 | 149 | 150 | 162 | 198 | 226 | 262 | 276 | 288 | 315 | 359 | 363 | 365 | 404 | 443 | 444 | 451 | 480 | 492 | 496 |
| **BNG924** | T | A | R | K | S | Q | I | Q | S | L | R | D | I | D | K | I | P | I | K | V | I | R | R | M | K | R | V |
| **BNG943** | S | A | Q | T | L | K | V | Q | G | Q | Q | N | V | N | K | V | H | V | K | V | L | K | M | L | R | K | I |
| **BNG948** | T | S | Q | T | S | Q | I | L | S | L | R | T | V | N | R | I | P | V | N | I | L | R | R | L | R | K | V |
| **Strain** | **Residue Position (NA)** |  |  |  |  |  |  |  |  |  |  |  |  |  |  |  |  |  |  |  |  |  |  |  |  |  |  |
|  | **22** | **34** | **54** | **60** | **64** | **75** | **123** | **129** | **137** | **192** | **237** | **245** | **246** | **290** | **294** | **310** | **340** | **342** | **362** | **367** | **381** |  |  |  |  |  |  |
| **BNG924** | N | Q | I | Y | A | Q | D | G | Y | L | I | A | H | V | V | N | D | V | I | V | E |  |  |  |  |  |  |
| **BNG943** | S | P | V | C | T | Q | N | D | H | M | L | V | H | A | I | D | D | I | I | V | E |  |  |  |  |  |  |
| **BNG948** | N | P | I | C | A | H | N | G | Y | L | I | I | R | A | I | N | N | V | V | A | G |  |  |  |  |  |  |
| **HA** | **BNG924** | **BNG943** | **BNG948** |  |  |  |  |  |  |  |  |  |  |  |  |  |  |  |  |  |  |  |  |  |  |  |  |
| **BNG924** | 0 |  |  |  |  |  |  |  |  |  |  |  |  |  |  |  |  |  |  |  |  |  |  |  |  |  |  |
| **BNG943** | 22 (4.06%) | 0 |  |  |  |  |  |  |  |  |  |  |  |  |  |  |  |  |  |  |  |  |  |  |  |  |  |
| **BNG948** | 15 (2.77%) | 18 (3.32%) | 0 |  |  |  |  |  |  |  |  |  |  |  |  |  |  |  |  |  |  |  |  |  |  |  |  |
| **NA** | **BNG924** | **BNG943** | **BNG948** |  |  |  |  |  |  |  |  |  |  |  |  |  |  |  |  |  |  |  |  |  |  |  |  |
| **BNG924** | 0 |  |  |  |  |  |  |  |  |  |  |  |  |  |  |  |  |  |  |  |  |  |  |  |  |  |  |
| **BNG943** | 15 (3.33%) | 0 |  |  |  |  |  |  |  |  |  |  |  |  |  |  |  |  |  |  |  |  |  |  |  |  |  |
| **BNG948** | 12 (2.66%) | 16 (3.55%) | 0 |  |  |  |  |  |  |  |  |  |  |  |  |  |  |  |  |  |  |  |  |  |  |  |  |

The representative viruses incorporate a total of 27 mutations, including all 22 mutations defining the monophyletic HA groups. Due to the variance of the orange monophyletic HA group, some of the unique mutations are included within this group’s representative strain (BNG924); the representative strain for the green monophyletic HA group (BNG943) contained 21 conserved mutations, 17 unique, yet the representative strain encompasses 13 unique mutations. Consequently, the orange monophyletic HA group (BNG924) has 10 unique mutations and yellow (BNG948) 5 mutations between the representative strains.

Of the amino acid substitutions within HA between the representative strains, only four are known antigenic epitopes, 149, 150, 198 and 226 (mature H9 HA numbering). Alternatively, non-antigenic external mutations 22-69 are situated at the base of the HA1 chain, 162, 259, 262, 276 and 288 sit within the head and cleavage hinge, 315 at start of the HA cleavage motif, 363 and 365 sit within the fusion peptide of the HA2 chain, 404, 443, 444 and 451 within the stem and fusion peptide, and 480, 492 and 496 at the anchor base.

Of the mutations within NA, only two of the amino acid variations are known antigenic epitope sites: 340 and 342 (H3N2 universal numbering = +18 amino acids [25]), forming part of the structural loops surrounding the enzymatic pocket. Interestingly, each representative virus has different combinations of amino acids at these two positions: BNG924 = DDV, BNG943 = DDI, BNG948 = NDV. The non-antigenic external mutations 22, 34, 54, 60, 64 and 75 form the flexible stalk and neck regions of NA and 123, 129, 137, 192, 237, 245, 246, 310, 362, 367 and 381 map to the internal structure.

The representative HAs varied from 2.77% (15 mutations) to 4.06% (22 mutations) within the mature sequence and the NAs from 2.66% (12 mutations) to 3.55% (15 mutations). Both HA and NA of BNG924 and BNG948 were most similar; alternatively, HA was most diverse between BNG924 and BNG943 and NA most diverse between BNG943 and BNG948 (3.33%), closely followed by BNG924 (3.55%).

N-linked glycans were predicted for both HA (n=8) and NA (n=8). Within HA, the number of sites remained identical with the only difference between the strains being the composition of the N-linked glycan motif (N-X-S/T) where site 287 varies from NISK (BNG924 and BNG948) to NVSK (BNG943). The substitution from I to V increases the N-linked glycan potential but both motifs maintain high potential scores (0.6997 and 0.7316).

More variation in NA was observed, namely BNG943 again presenting varied motif arrangements. Sites 54 and 131 (mature H9 numbering) vary from NGTI and NGTT to NGTV and NDTT, respectively, but maintains potentials. Additionally, there are some slight variations in the confidence of the sites between strains which would be attributed to up and downstream sequence mutations. Site 29 was shared by all three representative viruses but BNG943 displays a lower potential to a negative result (0.4762). The motif of site 71 was shared by all strains but BNG943 exhibited an increased potential (0.6147). Similarly, site 387 differs in potential for BNG948 (0.4450) but retains the same motif and remains a negative result for all viruses. Alternatively, position 131 within BNG943 boasts a considerable decrease in potential at this site (0.5804) and a variation in motif (NDTT). Of the five considerable substitutions in potential or motifs, four are exhibited by BNG943, two of which were alterations of the N-linked glycan motifs, and one by BNG948.

To assess antigenicity, recombinant viruses containing the HA and its corresponding NA of each reference strain, were propagated using reverse genetics (RG) combining remaining genes from the laboratory-adapted strain A/Puerto Rico/8/1934 (PR8). Whole virus antisera were raised to each strain in specific-pathogen free (SPF) White Leghorn chickens. The final blood draw (42-days post-primary vaccination) provided the antisera for serological characterisation and assessment of cross-reactivity by haemagglutination inhibition (HI) assay and virus neutralisation test (VNT) (Figure 5).

**Figure 5:**
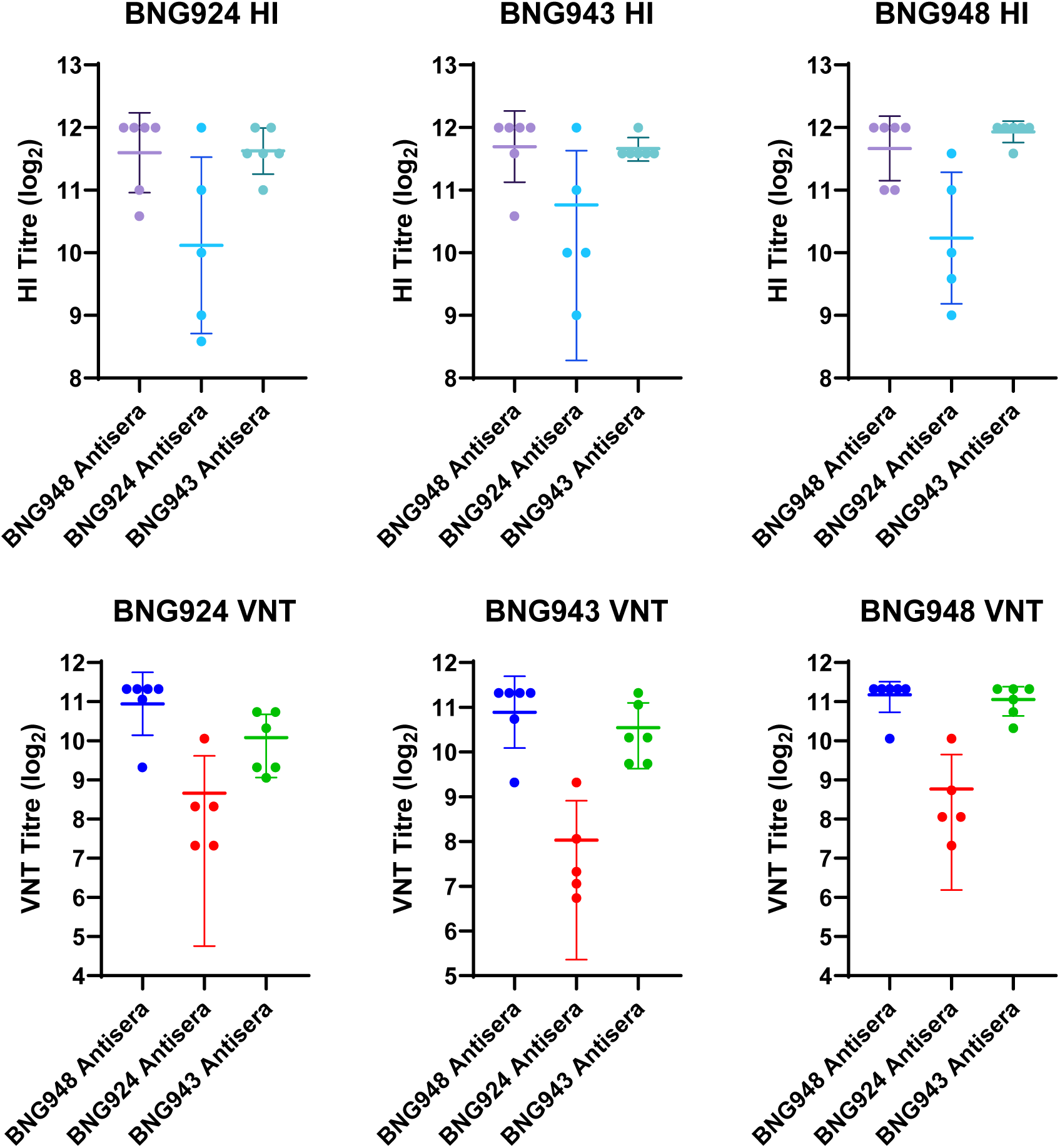
Haemagglutination inhibition (HI) and virus neutralisation test (VNT) titres of H9N2 viruses representing the monophyletic groups of HA. Each graph depicts the assay of the titled antigen. Each dot represents the geometric mean titre of an individual sera across three technical repeats with mean and standard error bars.

HI titres for each virus elicited high protective homologous and heterologous titres. Noticeably, BNG924 presented a reduced homologous immunogenicity, with a variation of 4 log_2_ HI titre across the biological repeats. The titres applied similarly to the heterologous viruses BNG943 and BNG948. However, heterologous HI titres to the BNG924 matched that of the heterologous sera, with a 1-1.5 log_2_ intra-sera variation. This suggests that while all three viruses are antigenically identical, BNG924 may demonstrate variable immunopotency, with only two of five birds responding comparatively to the heterologous sera. The same phenomenon was observed in the VNT titres; BNG924 elicited significantly lower protection titres to homologous and heterologous viruses, again with considerable intragroup range, with only one serum responding comparatively. Alternatively, BNG943 and BNG948 were highly cross-reactive to every representative virus, with minimal variations in VNT titres.

As the drop in immunopotency of BNG924 may be the result of distorted titration by HAU, each virus was alternatively titrated by plaque-forming units (pfu) and tissue culture infectious dose (TCID) (Table S5). All three representative viruses presented similar titres for all assays and similar plaque morphologies, eliminating the risk of unstable virus propagation of BNG924 and therefore vaccine seed.

These combined results can conclude that all three representative viruses for the co-circulating monophyletic groups of HA are structurally antigenically similar, but BNG924 exhibits a heterogeneous primary seroconversion pattern across vaccinated hosts despite remaining fully sensitive to heterologous functional neutralisation.

## Discussion

This study sought to genetically and antigenically characterise a panel of H9N2 AIVs sampled from poultry across regions within Bangladesh.

While previous studies published time-scaled phylodynamic analysis of the genetic divergence of H9N2 in Bangladesh, our single-timepoint cross-sectional sampling revealed the co-existence of three distinct monophyletic HA groups of the G-lineage [7, 27]. This suggests synergistic harmony without competition between the divergent strains or ecological or geographical segregation and independent evolution. Each monophyletic group was largely characterised by distinct amino acid residues with some unique to geographical regions. Publicly available H9Nx sequencing data of HA in Bangladesh from 2020-2026 similarly presented three distinct monophyletic groups of HA and confirmed the reference strains for each group used in this study remain representative. This further confirms continuous synergistic evolution of strains across Bangladesh.

Similarly, a study by Carnegie *et al*., found genetic divergence between samples isolated in Dhaka and Chattogram districts [3]. Moreover, the same study found instances where viruses from distinct monophyletic groups were simultaneously isolated from the same source which coincided with two sites, Site 3 and Site 16, which found isolates from each monophyletic group. The lowermost (green) monophyletic group was strictly isolated from Chattogram or Cumilla only, whereas the middle (orange) and uppermost (red) monophyletic branches were widely distributed across all site clusters. The co-circulation of monophyletic groups is likely attributed to trade, with the capital Dhaka driving the movement nationally.

Carnegie *et al*., 2023 investigated spatial spread and dispersal patterns of H9N2 in Bangladeshi poultry and identified that epidemiological connectivity between major cities, such as Dhaka and Chattogram, suggests localised genetic heterogeneity [3]. The data from this study mimicked the general trend of diversity across all isolates. Additionally, Carnegie *et al*., 2023 identified sporadic introductions of genetically distinct viruses into distant markets. Again, this study too found the isolation of strains from various monophyletic groups from a single site, suggesting trade/bird movement or strong epidemiological connections between markets [28].

Carnegie *et al*., 2023 also observed tree topological incongruence between HA and NA within the Bangladeshi H9N2 population [3] concluding that continuous co-infection of birds in high-density live bird markets creates an ideal environment for H9N2 variants to exchange HA and NA segments with one another without changing subtype. Within this study, we were able to extrapolate genetic data of HA and its paired NA from phylogenetic arrangement within the tanglegram. Using a continuous, quantitative composite genetic mismatch, we were able to measure degrees of genetic distance across segments and evaluate local neighbourhood conservation of fixed trees. From the 219 strains, 20 were predicted to be reassortants, of which 17 resided in the distinct green monophyletic group geographically localised in Chattogram and Cumilla. Reassortants of H9N2 HA and NA specifically have been identified in Bangladesh previously, with subtype maintenance yet reassortment between different endemic sub-lineages of NA carrying unique molecular markers resulting in various genotypes [27]. H9N2 is also highly known for its frequent reassortment of internal genes, namely PB2 and NS, with other subtypes, including H7N3 and H5N1, often resulting in highly fit reassortants [29, 30].

The four amino acid substitutions within HA identified as known antigenic epitopes, 149, 150, 198 and 226. 149, 150 and 198 are highly prone to positive selection pressure due to their proximity to the receptor binding site (RBS) [31–33]. 226 is another highly variable site on the globular head with mutations, particularly glycans, can shield critical neutralising epitopes [34]. Mutations 22-69 situated at the base of HA1 are key in anchoring and the structural framework of the stem [35] with limited variation and largely inaccessible by antibodies. 162, 259, 262, 276 and 288 sit within the head and cleavage hinge are structural and with the exception of 162, internalised and inaccessible to antibody binding [33, 36]. 315 is the start of the HA cleavage motif; proline (P) is most commonly present at this site and the variation to H fluctuates the motif from a tri- to a di-basic site which in turn allows the activation loop subtle increased physical elasticity. This mutation can suggest enhanced infectivity in the presence of secondary infections and antigen repertoires of the chicken respiratory tract [37]. 363 and 365 sit within the fusion peptide of the HA2 chain, 404, 443, 444 and 451 provide structural integrity of the stem and pre-fusion trimer, and 480, 492 and 496 at the anchor base, stabilising the transmembrane domain [35, 38, 39]. The genetic variation between the strain certainly suggests localised evolutionary drift, and as such monophyletic lineage grouping, with some mutations potentially modifying the virus to situational fitness adaptation.

NA mutations 340 and 342 surrounding the enzymatic pocket can modify activity as well as driving antigenic variation [40]. The variation at these sites between the strains can suggest that this may be an active site for antigenic drift. Alternatively, 22, 34, 54, 60, 64 and 75 form the flexible stalk and neck regions of NA where antibody binding does not neutralise virus activity. 123, 129, 137, 192, 237, 245, 246, 310, 362, 367 and 381 map to the internal strands of the core matrix and are therefore inaccessible to host antibodies [40].

The N-linked glycan potentials of each candidate HA were identical in position and likelihood but differ in motif at one position. Position 11 lies within the signal peptide and is cleaved during protein maturation so would not exist on the viral surface [36]. Position 123 is located within the major antigenic region, the receptor binding site, of the globular head and is a known marker for antigenic drift; the glycan position sterically blocks the main antigenic loops for neutralising antibodies [41]. Positions 280 and 287 are co-occurring glycan sites located towards the base of the HA head and may interfere with antibody access to the HA stalk or the lower lateral surfaces of the head domain [42]. Site 474, albeit lower potential, is likely conserved to preserve structure and stability during fusion activation [43]. As all three viruses share major glycan profiles of the HA, namely 87, 123, 280 and 287, their epitope shielding profiles are similar and are therefore unlikely to contribute to antigenic diversity between the strains.

However, the N-linked glycan prediction for NA differs slightly between viruses. Similarly, all NAs exhibit the same position but differ quite considerably in potential and motifs. The N2 sequences map to a natural stalk deletion of two amino acids, a common evolutionary trait in chicken-adapted G-lineage H9N2 AIVs and thus conserved across Bangladesh [44, 45]. Position 29 is situated at the boundary between the N-terminal and hydrophobic transmembrane domain at the base of the stalk [46]. Position 46 is a canonical universal site conserved across virtually all N2 NAs and is critical for early protein folding and stability in the endoplasmic reticulum [46]. Position 54 is often variable as it sits within the variable stalk domain and dictates the structural flexibility and host adaptation of the virus [47]. Position 71 again corresponds to a canonical universal site and acts as a structural pivot between the stalk and globular head [46]. Within the globular head sits the remaining four variable sites: 131, 185, 219 and 387 (canonical numbering: 133, 187, 221 and 392, respectively). Position 131 is a highly conserved site located near the top of the head and is vital in structurally stabilising the enzymatic pocket [48]. Position 185 is within the upper loop flanking the catalytic site and the loss or gain of a N-linked glycan through mutation typically expose or conceal critical antigenic epitopes [49]. Similarly, position 219 sits on another hyper-variable outer-loop of the globular head and primarily influences antigenic shielding and enzymatic activity [27]. Position 387 again is a conserved lateral-side site towards the base of the globular head and protects the integrity of the underlying structural scaffolding of the tetramer [50].

Despite considerable genetic diversity between the representative strains, the antigenic phenotype was remarkably retained. Previous publications have highlighted this paradox within Bangladesh of chicken-origin H9N2, with viruses from other hosts, such as quail, displaying antigenic drift [33, 51]. Other studies have suggested that rapid antigenic diversification in H9N2 occurs primarily in regions where inactivated poultry vaccines are widely deployed, encouraging immune escape mutants at key antigenic sites [31, 52]. Bangladesh doesn’t have a strict vaccination regime such as that in China, but some commercial vaccines are available, namely A/chicken/Bangladesh/NRL-H9-1767/2022 (G5.7/G1-h9.4.2 clade), used by Bangladesh Livestock Research Institute (BLRI) Local Strain Vaccine [53], which aligns with the green HA subgroup (Figure S1) and historical A/Quail/Hong Kong/G1/1997 (G-lineage) used by Incepta Pharmaceuticals (GuardFlu Vet) and Ceva (Cevac® Flu H9) [54], the reference strain for the lineage. Other vaccines covering the B-lineage (A/chicken/Beijing/1/1994) are also distributed by FnF Pharmaceuticals (Bangla Avianza Vac H9) again representing the ancestral lineage reference strain [44]. This may suggest that despite the high genetic variation and dated vaccine seed, the antigenic diversity within the lineage appears to remain limited despite numerous mutations in HA and NA and thus ancestral strains remain effective against significant disease but not spread [1].

In contrast, the reduced immunopotency seen by the most dominant and widespread strain BNG924 may challenge vaccination efficacy if used as a candidate vaccine seed. The reduced induction of immunity could be the result of low dosing or unstable virus within the vaccine, yet three methods of titrations and glycan presentation warranted highly similar values across all three representative viruses. This observation could be a disconnect between *in vivo* and *in vitro* potency, namely intracellular trafficking/endosomal escape, endogenous processing or defective particles [55]. In contrast, several studies have observed a similar disconnect between neutralisation cross-reactivity and immunogenicity, which may reflect differences in viral receptor-binding avidity and/or antibody-mediated blocking avidity, as well as antigen flexibility and modulation of innate immunity [56–59]. Mutations outside of traditional antigenic sites and the receptor binding domain may cause immunogenic asymmetry without directly interfering with the antigenic profile of the strain. Nonetheless, the substitutions identified between the strains can conclude to their lack of antigenic influence, and alternatively lack of immune potency, and aid antigen design for future vaccine seeds.

## Conclusion

H9N2 AIV continues to persist in Bangladeshi poultry. This study shows that despite considerable localised viral genetic variation across Bangladesh, antigenicity remains highly comparable between monophyletic groups of HA. Most notably was sporadic introductions of genetically variable strains were isolated from the same market suggesting a key epidemiological link or trading network between markets and could suggest a driver of antigenic shift and novel reassortants. Combined, these findings suggest that H9N2 in Bangladesh is continuously undergoing evolutionary change, despite retaining antigenicity.

Furthermore, our data suggested reduced protective efficacy of antibodies induced by one of the representative viruses, both dominant and nationally distributed. Despite TCID50, pfu and HAU being comparable between the representative strains, further evaluation and repeated *in vivo* analysis is required to conclude the reduced immunopotency observed. If so, strains which induce poor immunity may give rise to antigenic or phenotypic variants able to escape previous immunity or currently adopted vaccines for poultry.

## Methods

### Whole genome sequencing and assembly

Positive isolates taken from poultry across Bangladesh were subjected for whole genome sequencing at the Animal Plant and Health Agency (APHA) as previously described [60]. Influenza virus genomes were assembled from raw sequence data with the iterative refinement meta-assembler (IRMA) pipeline [61].

### Phylogenetic analysis

Whole haemagglutinin (HA) and neuraminidase (NA) amino acid sequences were aligned and trimmed to the mature open-reading frame (ORF), removing the signal peptide and cytoplasmic regions using MEGA-X [26]. Mid-point rooted maximum-likelihood trees constructed with the General Time Reversible model with discrete Gamma distribution and invariant sites (GTR + G + I) with synapomorphic amino acid mutations that are present in sequential branches labelled at the nodes, were generated in RStudio [15] using ape [16], phangorn [17], ggtree [18] & tidyverse [19] packages.

A co-phyletic tanglegram was also generated in RStudio, utilising the amino acid phylogenetic trees, connecting the HA and NAs of the same strain, with ape [16] and phytools [20] packages. Connecting lines were coloured according to the monophyletic groups of HA.

Utilising the tanglegram, a dual-metric threshold system combining composite genetic mismatch scores with k-nearest neighbour topological overlaps. Composite genetic mismatch score was calculated by averaging normalised procrustes residuals and mean patristic distance discordances across cophenetic tree spaces. Higher scores denote strains with profound HA-NA topological decoupling. The k-nearest neighbour overlap was calculated by identifying the ten closest genetic relatives (k=10) for each isolate based on patristic distance in the HA tree and determining the proportion of those same relatives retained among its ten closest neighbours in the NA tree. K=10 was selected to capture local outbreak cluster dynamics while avoiding sensitivity to minor phylogenetic node order instability among closely related isolates. This threshold effectively defines the immediate genetic clade background without incorporating broader macro-population tree topology.

Strains were categorised into three distinct evolutionary states: true reassortants (mismatch score > Mean + 1.5 standard deviation (StD) and neighbour overlap ≤ 30%) representing inter-lineage segment exchange; co-evolved/variants (neighbour overlap ≥ 60%), representing linked HA-NA inheritance with lineage-specific drift; and intermediate/drift isolates (neighbour overlap 31-59%), reflecting intra-clade micro-evolutionary variation.

### Virus selection

One virus per monophyletic branch was selected as the candidate for that subgroup of the HA G-lineage. A consensus was made of each monophyletic branch and the closest isolate to that consensus selected.

### Generation of viruses through reverse genetics (RG)

Viruses were generated using reverse genetics (RG) using the eight-segment bi-directional plasmid system [62]. Representative viruses of the subgroups were generated as haemagglutinin (HA) and neuraminidase (NA) homologous genes with the remaining six core internal genes from of A/Puerto Rico/8/1934 (PR8) H1N1 virus.

For virus rescue, human embryonated kidney (HEK-293T) cells were cultured in T25 flasks until they reached 90-100% confluence and maintained in Dulbecco’s Modified Eagle Medium) (DMEM) supplemented with 10% foetal calf serum (FCS) and 1% penicillin/streptomycin. For transfection, cells were washed with phosphate-buffered saline (PBS) and a plasmid mixture containing 1µg of each plasmid was diluted in OptiMem™ and incubated with Lipofectamine™ for 30 minutes before being added to the cells for 16 hours. Following transfection, the cells were washed in PBS and incubated in DMEM with tosyl phenylalanyl chloromethyl ketone (TPCK)-treated trypsin for 72 hours, while monitoring for cytopathic effect (CPE). Cell culture supernatants were harvested, and successful virus rescue was confirmed by haemagglutination assay. Recovered viruses were subsequently propagated in 10 day-old embryonated chicken eggs (VALO BioMedia, GmbH).

### Haemagglutination assay and Haemagglutination inhibition (HI) assay

Haemagglutination and haemagglutination inhibition (HI) assays were performed to determine haemagglutination titre of viruses propagated in embryonated chicken eggs and to quantify antibodies in serum from vaccinated birds capable of inhibiting agglutination of chicken red blood cells (RBCs), respectively. Briefly, freshly prepared (in-house) chicken RBCs were washed three times in phosphate-buffered saline (PBS) and diluted to a working concentration of 1% (v/v) in PBS.

For haemagglutination titration, 50µL of virus was serially diluted two-fold in 50µL of PBS in 96-well v-bottom plates. An aliquot of 50µL of chilled 1% RBC suspension were added to each well and incubated at 4°C for 45-60 minutes. Plates were tilted to observe for agglutination or absence of, confirmed by a teardrop shape formed from pelleted RBCs. Virus titres were expressed as log_2_ haemagglutinating units (HAU).

A standardised dose of 4 HAU of each virus was used for HI assays. Briefly, 25µL of homologous antisera raised against the corresponding non-mutated virus were serially two-fold diluted in duplicate with 25µL of PBS in 96-well v-bottom plates. Subsequently, 25µL of virus (4 HAU) was added to each well and incubated at 37°C for 60 minutes. Following incubation, 50µL of chilled 1% RBC suspension was added to each well, and the plates were further incubated at 4°C for 45-60 minutes. HI titres were recorded as the reciprocal of the highest serum dilution that completely inhibited haemagglutination as indicated by the formation of a distinct teardrop-shaped RBC pellet upon tilting the plate. Titres were expressed as log₂ HI titres.

### Raising antisera

Recombinant virus stocks were propagated in embryonated chicken eggs and subsequently inactivated with 0.1% (v/v) β-propiolactone (BPL) (Sigma-Aldrich). Complete virus inactivation was verified by inoculating 200µL of each inactivated virus preparation into embryonated eggs over three sequential passages using allantoic fluid from the preceding passage for each subsequent inoculation. Following each passage, viral replication was assessed by haemagglutination assay. Any virus preparation exhibiting detectable HA activity was subjected to a further round of BPL inactivation and re-tested to confirm complete inactivation.

Inactivated virus preparations were concentrated via ultracentrifugation and formulated into vaccines containing 1024 HAU per dose (dosing supported by Kilany *et al*., 2016 [63]), calculated by haemagglutination assay post-inactivation across three technical repeats. Vaccines were prepared at a 30:70 antigen-to-adjuvant ratio using MONTANIDE™ ISA 71 VG adjuvant (SEPPIC), according to the manufacturer’s instructions.

Specific pathogen-free (SPF) day-old White Leghorn chickens were vaccinated subcutaneously with 200µL of vaccine and received a booster vaccination 14 days after the primary immunisation. Blood samples were collected at 42 days post-vaccination marking the end of the experiment. Serum was separated from whole blood and stored at −20°C until use in HI and VNT assays.

### Tissue culture infectious dose (TCID_50_) and virus neutralisation test (VNT)

TCID50 assay determined recombinant virus titres for subsequent VNT assays; a standardised in-house H9N2 G-lineage recombinant virus (A/Chicken/Pakistan/UDL01/2008 (UDL)) was used as a positive assay control. Confluent MDCK cells seeded in 96-well plates were washed with PBS and infected with two-fold serial dilution of virus in quadruplicate for 1 hour. Following adsorption, the inoculum was removed and replaced with DMEM supplemented with TPCK-treated trypsin. Cells were then incubated for 72 hours at 37°C. After incubation, the cells monolayers were stained with crystal violet solution and viral titre were calculated using the Spearman-Karber method:

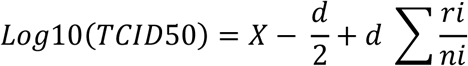

Where *X* is the lowest dilution where all wells were positive for viral infection, *d* is the log_10_ of the dilution factor (here 0.5), *ri* is the number of positive wells and *ni* is the number of replicates.

VNT assays were performed in triplicate using confluent monolayers of MDCK cells prepared as described above. Chicken antisera obtained from vaccinated birds were heat-inactivated at 56°C for 30 minutes prior to use; similarly, UDL with homologous antisera was used as a positive assay control. Two-fold serial dilutions of each antiserum were prepared in DMEM and mixed with an equal volume of virus containing 150 TCID50 of the respective virus per well in separate 96-well plates. Virus only controls to ensure infective dose titres were incubated without antisera. The virus-antiserum mixtures were incubated at 37°C for 1 hour to allow antibody binding and neutralisation. Meanwhile, MDCK cell monolayers were washed with phosphate-buffered saline (PBS). Following incubation, the virus-antiserum mixture was added to the cells and incubated at 37°C for further 72 hours. Cells were then stained with crystal violet solution and examined for the presence or absence of virus-induced cytopathic effects (CPE). VNT titres were defined as the reciprocal of the highest serum dilution that completely inhibited CPE in the infected cell monolayer.

### Plaque assay

Plaque assays were performed in duplicate to alternatively titrate representative viruses. MDCK cells were seeded in 12-well plates for next day confluency. Viruses were serially diluted 10-fold and incubated in growth medium lacking FCS at 37°C and 5% CO₂ for 1 hour, with each dilution occupying two wells. The virus medium was removed, and cells washed prior to coating with cooled molten 2% agarose agar combined with overlay media (MEM (Gibco), 0.2% w/v filtered bovine serum albumin, 200nM L-glutamine in saline (Sigma), 7.5% NaCO_2_, 10M Hepes, 1% dextran hydrochloride (Sigma), 1% penicillin-streptomycin (Gibco)) supplemented with TPCK at 1.0μg/ml. Plates were stored upside-down at 37°C and 5% CO₂ for a further 72 hours. Cells were fixed and stained with crystal violet stain (70% distilled water, 20% methanol, 10% 1% crystal violet in water).

## Ethics approval

All animal studies and procedures were conducted in accordance with European and United Kingdom Home Office regulations, including the Animals (Scientific Procedures) Act 1986 Amendment Regulations 2012, under approved project licenses (P68D44CF4 and PP6471846) issued by the UK Home Office. All experimental procedures were reviewed and approved by the Animal Welfare and Ethical Review Board at The Pirbright Institute and were performed by highly trained and licensed personnel. Animal welfare was closely monitored throughout the studies, with adherence to the 3Rs principles wherever possible.

## Data availability

All sequence data analysed in this study are publicly available through the GISAID Initiative and the NCBI Nucleotide database. Raw experimental datasets, plasmids, and viral isolates are available from the corresponding author upon reasonable request and in accordance with institutional policies and applicable biosafety regulations.

## Acknowledgements

We gratefully acknowledge the collaborative contributions of the Bangladesh Livestock Research Institute (BLRI), Chattogram Veterinary and Animal Sciences University (CVASU), The Royal Veterinary College (RVC), The Animal and Plant Health Agency (APHA), and The Pirbright Institute. We thank the staff of the Avian Influenza and Newcastle Disease Laboratory and Animal Services at The Pirbright Institute for their invaluable support in facilitating this research and generating the *in vivo* samples used in this study.

We also gratefully acknowledge all data contributors, namely the authors and originating laboratories responsible for obtaining the specimens, and the submitting laboratories for generating and sharing genetic sequence data and associated metadata through the GISAID Initiative, on which this work is based.

## Author Contributions

|  |  |
|---|---|
| Rebecca Daines (RD) | Data curation, formal analysis, methodology, validation, publication drafting |
| Jean-Remy Sadeyen (J-RS) | Resources, data curation |
| Mahmudul Hasan (MH) | Resources, data curation |
| Tridip Das (TD) | Resources, data curation |
| Rashed Mahmud Sizer (RMS) | Resources, data curation |
| Md Helal Uddin (MHU) | Resources, data curation |
| Paritosh Kumar Biswas (PKB) | Resources, data curation |
| Md Abdus Samad (MAS) | Resources, data curation |
| Md Ahasanul Hoque (MAH) | Resources, data curation |
| Anne Conan (AC) | Resources, data curation |
| Joshua Lynton-Jenkins (JL-J) | Data curation, formal analysis |
| Ashley Banyard (AB) | Conceptualisation, project administration |
| Ian Brown (IB) | Conceptualisation, publication review and editing |
| Tom Lewis (TL) | Resources, data curation, formal analysis |
| Jayna Raghwani (JR) | Resources, data curation, formal analysis |
| Guillaume Fournie (GF) | Formal analysis, project administration, conceptualisation |
| Damer Blake (DB) | Funding acquisition, project administration, conceptualisation |
| Fiona Tomley (FT) | Funding acquisition, project administration, conceptualisation, publication review and editing |
| Munir Iqbal (MI) | Funding acquisition, project administration, conceptualisation, publication review and editing |

## Supplementary Tables and Figures

**Table S1:**
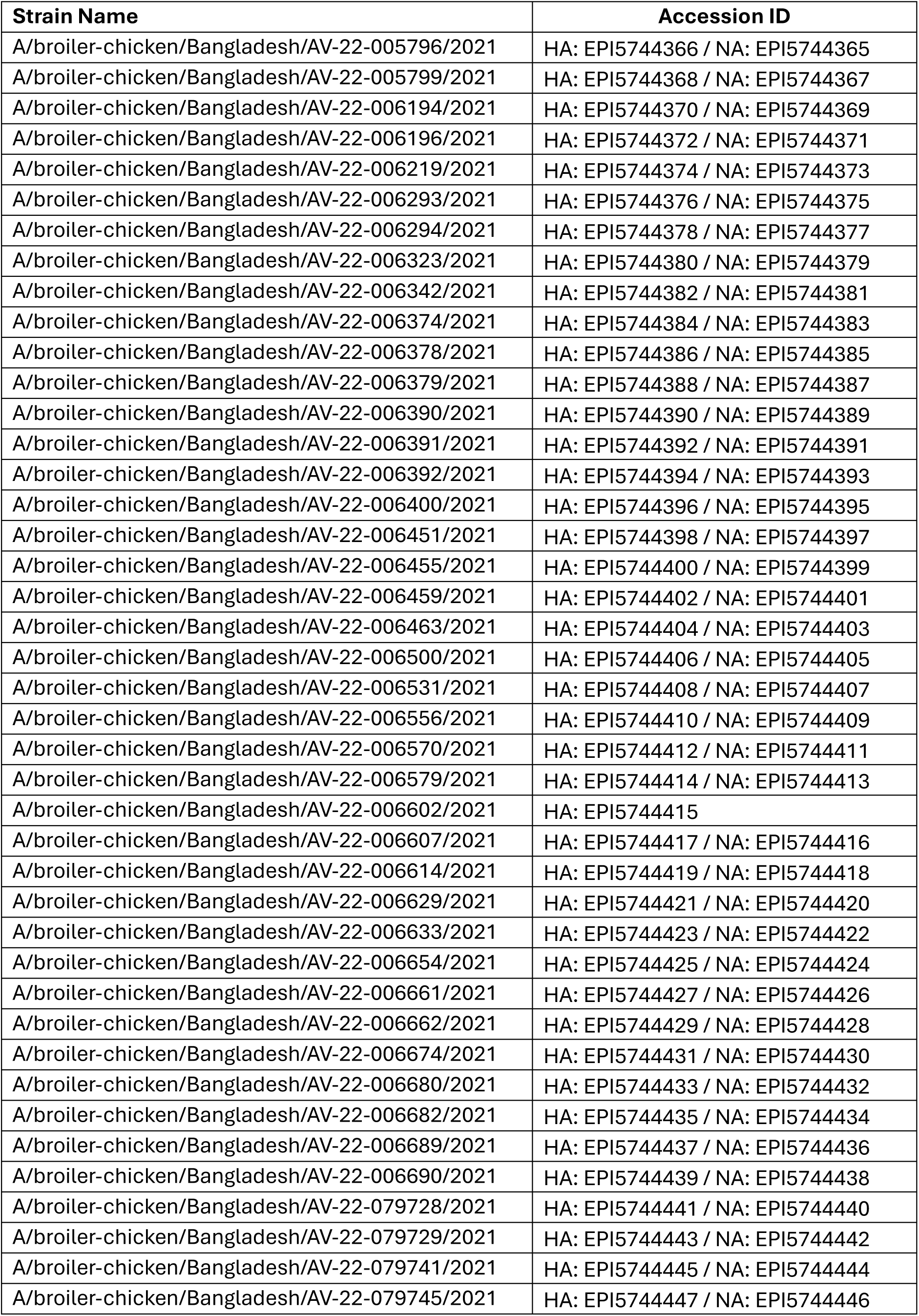
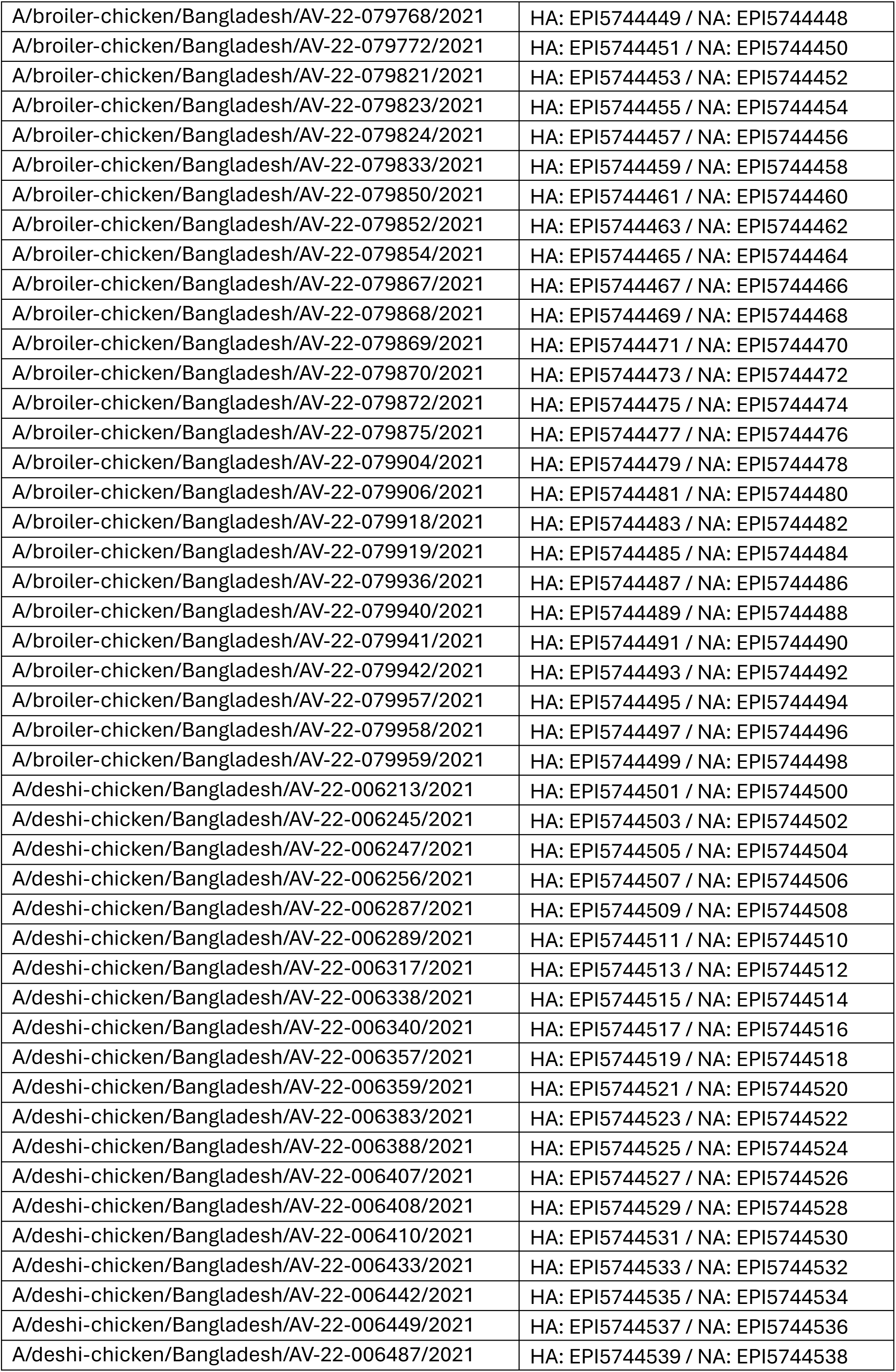
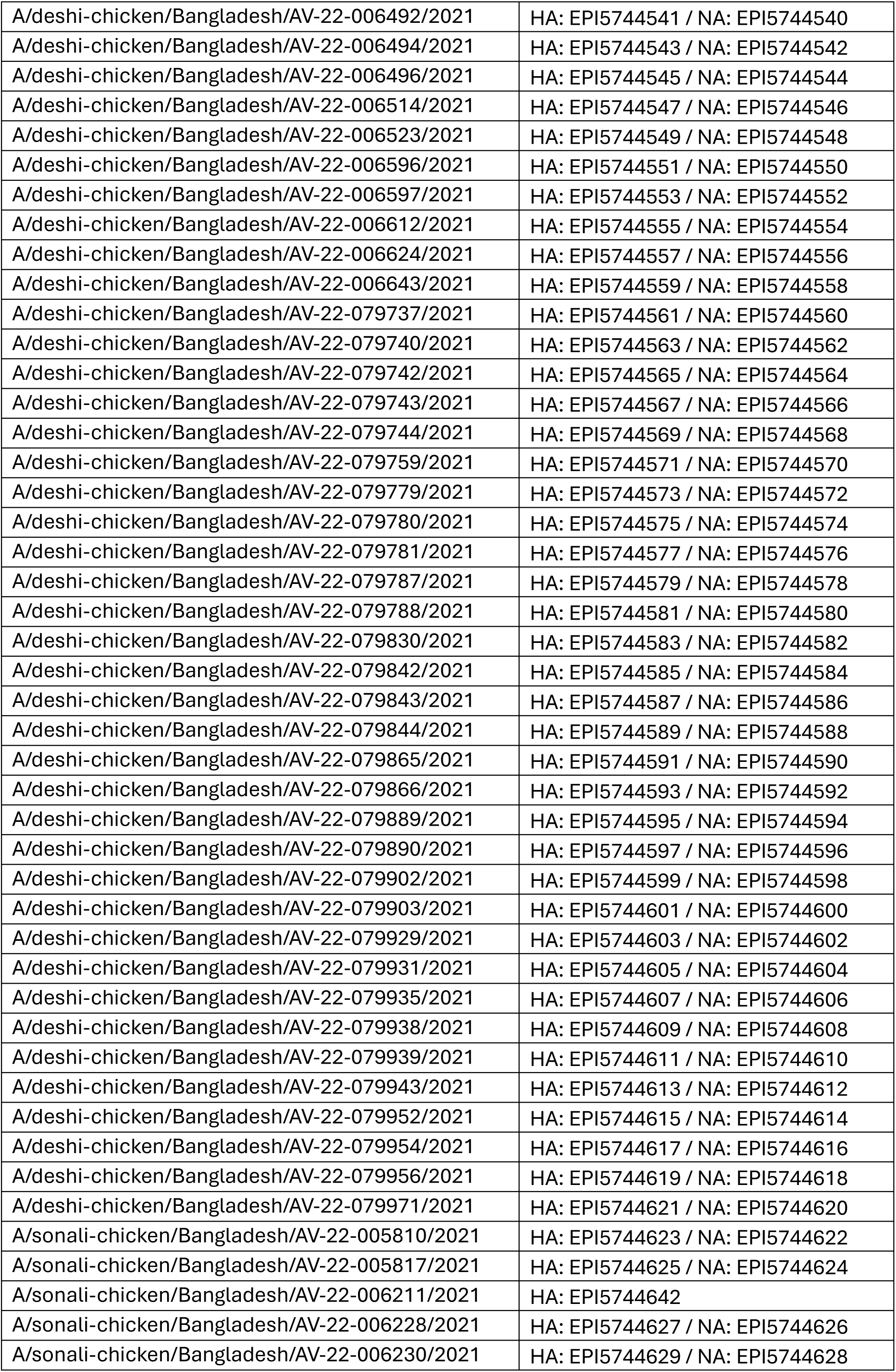
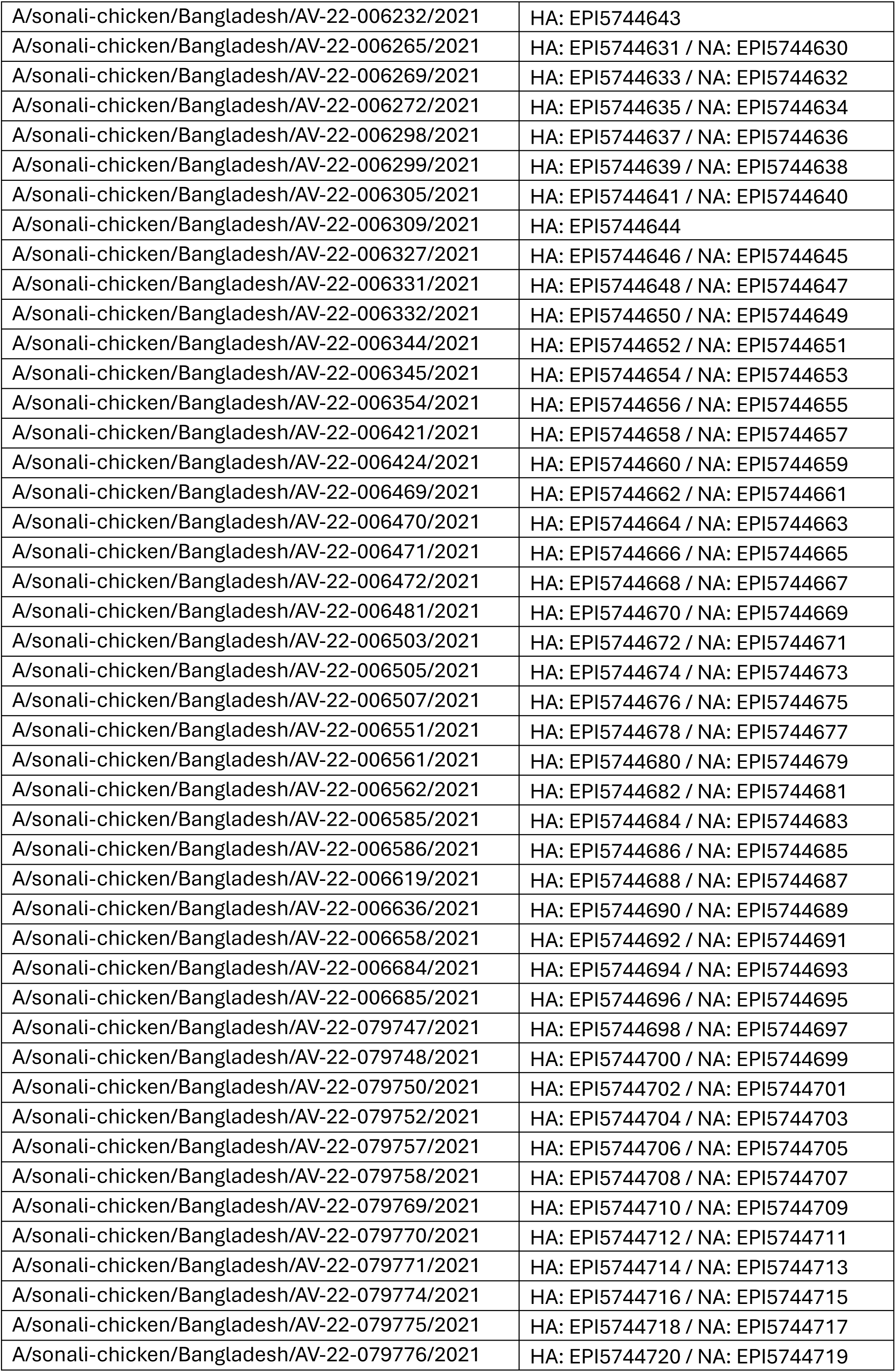
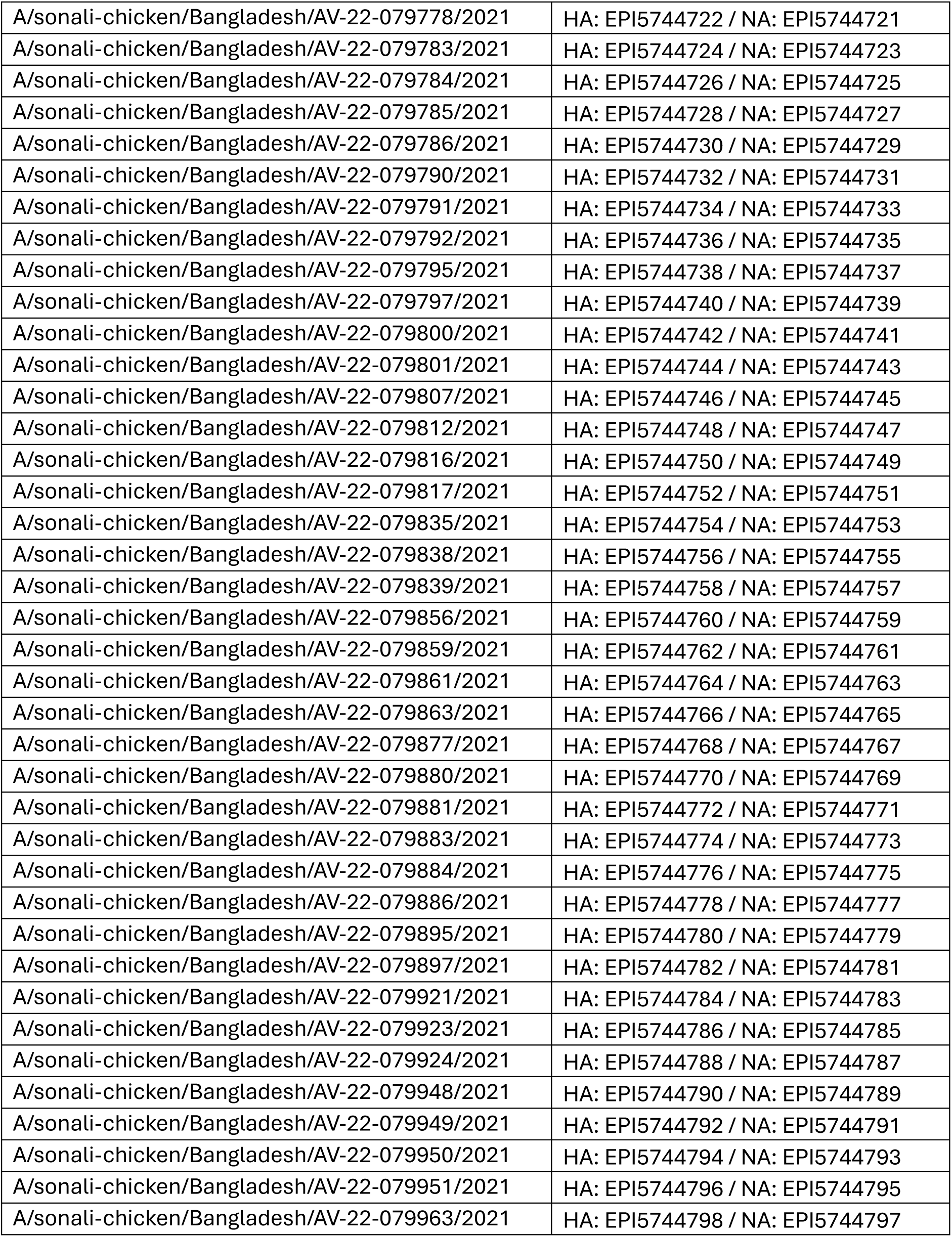
Isolate accession numbers (GISAID).

| Strain Name | Accession ID |
|---|---|
| A/broiler-chicken/Bangladesh/AV-22-005796/2021 | HA: EPI5744366 / NA: EPI5744365 |
| A/broiler-chicken/Bangladesh/AV-22-005799/2021 | HA: EPI5744368 / NA: EPI5744367 |
| A/broiler-chicken/Bangladesh/AV-22-006194/2021 | HA: EPI5744370 / NA: EPI5744369 |
| A/broiler-chicken/Bangladesh/AV-22-006196/2021 | HA: EPI5744372 / NA: EPI5744371 |
| A/broiler-chicken/Bangladesh/AV-22-006219/2021 | HA: EPI5744374 / NA: EPI5744373 |
| A/broiler-chicken/Bangladesh/AV-22-006293/2021 | HA: EPI5744376 / NA: EPI5744375 |
| A/broiler-chicken/Bangladesh/AV-22-006294/2021 | HA: EPI5744378 / NA: EPI5744377 |
| A/broiler-chicken/Bangladesh/AV-22-006323/2021 | HA: EPI5744380 / NA: EPI5744379 |
| A/broiler-chicken/Bangladesh/AV-22-006342/2021 | HA: EPI5744382 / NA: EPI5744381 |
| A/broiler-chicken/Bangladesh/AV-22-006374/2021 | HA: EPI5744384 / NA: EPI5744383 |
| A/broiler-chicken/Bangladesh/AV-22-006378/2021 | HA: EPI5744386 / NA: EPI5744385 |
| A/broiler-chicken/Bangladesh/AV-22-006379/2021 | HA: EPI5744388 / NA: EPI5744387 |
| A/broiler-chicken/Bangladesh/AV-22-006390/2021 | HA: EPI5744390 / NA: EPI5744389 |
| A/broiler-chicken/Bangladesh/AV-22-006391/2021 | HA: EPI5744392 / NA: EPI5744391 |
| A/broiler-chicken/Bangladesh/AV-22-006392/2021 | HA: EPI5744394 / NA: EPI5744393 |
| A/broiler-chicken/Bangladesh/AV-22-006400/2021 | HA: EPI5744396 / NA: EPI5744395 |
| A/broiler-chicken/Bangladesh/AV-22-006451/2021 | HA: EPI5744398 / NA: EPI5744397 |
| A/broiler-chicken/Bangladesh/AV-22-006455/2021 | HA: EPI5744400 / NA: EPI5744399 |
| A/broiler-chicken/Bangladesh/AV-22-006459/2021 | HA: EPI5744402 / NA: EPI5744401 |
| A/broiler-chicken/Bangladesh/AV-22-006463/2021 | HA: EPI5744404 / NA: EPI5744403 |
| A/broiler-chicken/Bangladesh/AV-22-006500/2021 | HA: EPI5744406 / NA: EPI5744405 |
| A/broiler-chicken/Bangladesh/AV-22-006531/2021 | HA: EPI5744408 / NA: EPI5744407 |
| A/broiler-chicken/Bangladesh/AV-22-006556/2021 | HA: EPI5744410 / NA: EPI5744409 |
| A/broiler-chicken/Bangladesh/AV-22-006570/2021 | HA: EPI5744412 / NA: EPI5744411 |
| A/broiler-chicken/Bangladesh/AV-22-006579/2021 | HA: EPI5744414 / NA: EPI5744413 |
| A/broiler-chicken/Bangladesh/AV-22-006602/2021 | HA: EPI5744415 |
| A/broiler-chicken/Bangladesh/AV-22-006607/2021 | HA: EPI5744417 / NA: EPI5744416 |
| A/broiler-chicken/Bangladesh/AV-22-006614/2021 | HA: EPI5744419 / NA: EPI5744418 |
| A/broiler-chicken/Bangladesh/AV-22-006629/2021 | HA: EPI5744421 / NA: EPI5744420 |
| A/broiler-chicken/Bangladesh/AV-22-006633/2021 | HA: EPI5744423 / NA: EPI5744422 |
| A/broiler-chicken/Bangladesh/AV-22-006654/2021 | HA: EPI5744425 / NA: EPI5744424 |
| A/broiler-chicken/Bangladesh/AV-22-006661/2021 | HA: EPI5744427 / NA: EPI5744426 |
| A/broiler-chicken/Bangladesh/AV-22-006662/2021 | HA: EPI5744429 / NA: EPI5744428 |
| A/broiler-chicken/Bangladesh/AV-22-006674/2021 | HA: EPI5744431 / NA: EPI5744430 |
| A/broiler-chicken/Bangladesh/AV-22-006680/2021 | HA: EPI5744433 / NA: EPI5744432 |
| A/broiler-chicken/Bangladesh/AV-22-006682/2021 | HA: EPI5744435 / NA: EPI5744434 |
| A/broiler-chicken/Bangladesh/AV-22-006689/2021 | HA: EPI5744437 / NA: EPI5744436 |
| A/broiler-chicken/Bangladesh/AV-22-006690/2021 | HA: EPI5744439 / NA: EPI5744438 |
| A/broiler-chicken/Bangladesh/AV-22-079728/2021 | HA: EPI5744441 / NA: EPI5744440 |
| A/broiler-chicken/Bangladesh/AV-22-079729/2021 | HA: EPI5744443 / NA: EPI5744442 |
| A/broiler-chicken/Bangladesh/AV-22-079741/2021 | HA: EPI5744445 / NA: EPI5744444 |
| A/broiler-chicken/Bangladesh/AV-22-079745/2021 | HA: EPI5744447 / NA: EPI5744446 |
| A/broiler-chicken/Bangladesh/AV-22-079768/2021 | HA: EPI5744449 / NA: EPI5744448 |
| A/broiler-chicken/Bangladesh/AV-22-079772/2021 | HA: EPI5744451 / NA: EPI5744450 |
| A/broiler-chicken/Bangladesh/AV-22-079821/2021 | HA: EPI5744453 / NA: EPI5744452 |
| A/broiler-chicken/Bangladesh/AV-22-079823/2021 | HA: EPI5744455 / NA: EPI5744454 |
| A/broiler-chicken/Bangladesh/AV-22-079824/2021 | HA: EPI5744457 / NA: EPI5744456 |
| A/broiler-chicken/Bangladesh/AV-22-079833/2021 | HA: EPI5744459 / NA: EPI5744458 |
| A/broiler-chicken/Bangladesh/AV-22-079850/2021 | HA: EPI5744461 / NA: EPI5744460 |
| A/broiler-chicken/Bangladesh/AV-22-079852/2021 | HA: EPI5744463 / NA: EPI5744462 |
| A/broiler-chicken/Bangladesh/AV-22-079854/2021 | HA: EPI5744465 / NA: EPI5744464 |
| A/broiler-chicken/Bangladesh/AV-22-079867/2021 | HA: EPI5744467 / NA: EPI5744466 |
| A/broiler-chicken/Bangladesh/AV-22-079868/2021 | HA: EPI5744469 / NA: EPI5744468 |
| A/broiler-chicken/Bangladesh/AV-22-079869/2021 | HA: EPI5744471 / NA: EPI5744470 |
| A/broiler-chicken/Bangladesh/AV-22-079870/2021 | HA: EPI5744473 / NA: EPI5744472 |
| A/broiler-chicken/Bangladesh/AV-22-079872/2021 | HA: EPI5744475 / NA: EPI5744474 |
| A/broiler-chicken/Bangladesh/AV-22-079875/2021 | HA: EPI5744477 / NA: EPI5744476 |
| A/broiler-chicken/Bangladesh/AV-22-079904/2021 | HA: EPI5744479 / NA: EPI5744478 |
| A/broiler-chicken/Bangladesh/AV-22-079906/2021 | HA: EPI5744481 / NA: EPI5744480 |
| A/broiler-chicken/Bangladesh/AV-22-079918/2021 | HA: EPI5744483 / NA: EPI5744482 |
| A/broiler-chicken/Bangladesh/AV-22-079919/2021 | HA: EPI5744485 / NA: EPI5744484 |
| A/broiler-chicken/Bangladesh/AV-22-079936/2021 | HA: EPI5744487 / NA: EPI5744486 |
| A/broiler-chicken/Bangladesh/AV-22-079940/2021 | HA: EPI5744489 / NA: EPI5744488 |
| A/broiler-chicken/Bangladesh/AV-22-079941/2021 | HA: EPI5744491 / NA: EPI5744490 |
| A/broiler-chicken/Bangladesh/AV-22-079942/2021 | HA: EPI5744493 / NA: EPI5744492 |
| A/broiler-chicken/Bangladesh/AV-22-079957/2021 | HA: EPI5744495 / NA: EPI5744494 |
| A/broiler-chicken/Bangladesh/AV-22-079958/2021 | HA: EPI5744497 / NA: EPI5744496 |
| A/broiler-chicken/Bangladesh/AV-22-079959/2021 | HA: EPI5744499 / NA: EPI5744498 |
| A/deshi-chicken/Bangladesh/AV-22-006213/2021 | HA: EPI5744501 / NA: EPI5744500 |
| A/deshi-chicken/Bangladesh/AV-22-006245/2021 | HA: EPI5744503 / NA: EPI5744502 |
| A/deshi-chicken/Bangladesh/AV-22-006247/2021 | HA: EPI5744505 / NA: EPI5744504 |
| A/deshi-chicken/Bangladesh/AV-22-006256/2021 | HA: EPI5744507 / NA: EPI5744506 |
| A/deshi-chicken/Bangladesh/AV-22-006287/2021 | HA: EPI5744509 / NA: EPI5744508 |
| A/deshi-chicken/Bangladesh/AV-22-006289/2021 | HA: EPI5744511 / NA: EPI5744510 |
| A/deshi-chicken/Bangladesh/AV-22-006317/2021 | HA: EPI5744513 / NA: EPI5744512 |
| A/deshi-chicken/Bangladesh/AV-22-006338/2021 | HA: EPI5744515 / NA: EPI5744514 |
| A/deshi-chicken/Bangladesh/AV-22-006340/2021 | HA: EPI5744517 / NA: EPI5744516 |
| A/deshi-chicken/Bangladesh/AV-22-006357/2021 | HA: EPI5744519 / NA: EPI5744518 |
| A/deshi-chicken/Bangladesh/AV-22-006359/2021 | HA: EPI5744521 / NA: EPI5744520 |
| A/deshi-chicken/Bangladesh/AV-22-006383/2021 | HA: EPI5744523 / NA: EPI5744522 |
| A/deshi-chicken/Bangladesh/AV-22-006388/2021 | HA: EPI5744525 / NA: EPI5744524 |
| A/deshi-chicken/Bangladesh/AV-22-006407/2021 | HA: EPI5744527 / NA: EPI5744526 |
| A/deshi-chicken/Bangladesh/AV-22-006408/2021 | HA: EPI5744529 / NA: EPI5744528 |
| A/deshi-chicken/Bangladesh/AV-22-006410/2021 | HA: EPI5744531 / NA: EPI5744530 |
| A/deshi-chicken/Bangladesh/AV-22-006433/2021 | HA: EPI5744533 / NA: EPI5744532 |
| A/deshi-chicken/Bangladesh/AV-22-006442/2021 | HA: EPI5744535 / NA: EPI5744534 |
| A/deshi-chicken/Bangladesh/AV-22-006449/2021 | HA: EPI5744537 / NA: EPI5744536 |
| A/deshi-chicken/Bangladesh/AV-22-006487/2021 | HA: EPI5744539 / NA: EPI5744538 |
| A/deshi-chicken/Bangladesh/AV-22-006492/2021 | HA: EPI5744541 / NA: EPI5744540 |
| A/deshi-chicken/Bangladesh/AV-22-006494/2021 | HA: EPI5744543 / NA: EPI5744542 |
| A/deshi-chicken/Bangladesh/AV-22-006496/2021 | HA: EPI5744545 / NA: EPI5744544 |
| A/deshi-chicken/Bangladesh/AV-22-006514/2021 | HA: EPI5744547 / NA: EPI5744546 |
| A/deshi-chicken/Bangladesh/AV-22-006523/2021 | HA: EPI5744549 / NA: EPI5744548 |
| A/deshi-chicken/Bangladesh/AV-22-006596/2021 | HA: EPI5744551 / NA: EPI5744550 |
| A/deshi-chicken/Bangladesh/AV-22-006597/2021 | HA: EPI5744553 / NA: EPI5744552 |
| A/deshi-chicken/Bangladesh/AV-22-006612/2021 | HA: EPI5744555 / NA: EPI5744554 |
| A/deshi-chicken/Bangladesh/AV-22-006624/2021 | HA: EPI5744557 / NA: EPI5744556 |
| A/deshi-chicken/Bangladesh/AV-22-006643/2021 | HA: EPI5744559 / NA: EPI5744558 |
| A/deshi-chicken/Bangladesh/AV-22-079737/2021 | HA: EPI5744561 / NA: EPI5744560 |
| A/deshi-chicken/Bangladesh/AV-22-079740/2021 | HA: EPI5744563 / NA: EPI5744562 |
| A/deshi-chicken/Bangladesh/AV-22-079742/2021 | HA: EPI5744565 / NA: EPI5744564 |
| A/deshi-chicken/Bangladesh/AV-22-079743/2021 | HA: EPI5744567 / NA: EPI5744566 |
| A/deshi-chicken/Bangladesh/AV-22-079744/2021 | HA: EPI5744569 / NA: EPI5744568 |
| A/deshi-chicken/Bangladesh/AV-22-079759/2021 | HA: EPI5744571 / NA: EPI5744570 |
| A/deshi-chicken/Bangladesh/AV-22-079779/2021 | HA: EPI5744573 / NA: EPI5744572 |
| A/deshi-chicken/Bangladesh/AV-22-079780/2021 | HA: EPI5744575 / NA: EPI5744574 |
| A/deshi-chicken/Bangladesh/AV-22-079781/2021 | HA: EPI5744577 / NA: EPI5744576 |
| A/deshi-chicken/Bangladesh/AV-22-079787/2021 | HA: EPI5744579 / NA: EPI5744578 |
| A/deshi-chicken/Bangladesh/AV-22-079788/2021 | HA: EPI5744581 / NA: EPI5744580 |
| A/deshi-chicken/Bangladesh/AV-22-079830/2021 | HA: EPI5744583 / NA: EPI5744582 |
| A/deshi-chicken/Bangladesh/AV-22-079842/2021 | HA: EPI5744585 / NA: EPI5744584 |
| A/deshi-chicken/Bangladesh/AV-22-079843/2021 | HA: EPI5744587 / NA: EPI5744586 |
| A/deshi-chicken/Bangladesh/AV-22-079844/2021 | HA: EPI5744589 / NA: EPI5744588 |
| A/deshi-chicken/Bangladesh/AV-22-079865/2021 | HA: EPI5744591 / NA: EPI5744590 |
| A/deshi-chicken/Bangladesh/AV-22-079866/2021 | HA: EPI5744593 / NA: EPI5744592 |
| A/deshi-chicken/Bangladesh/AV-22-079889/2021 | HA: EPI5744595 / NA: EPI5744594 |
| A/deshi-chicken/Bangladesh/AV-22-079890/2021 | HA: EPI5744597 / NA: EPI5744596 |
| A/deshi-chicken/Bangladesh/AV-22-079902/2021 | HA: EPI5744599 / NA: EPI5744598 |
| A/deshi-chicken/Bangladesh/AV-22-079903/2021 | HA: EPI5744601 / NA: EPI5744600 |
| A/deshi-chicken/Bangladesh/AV-22-079929/2021 | HA: EPI5744603 / NA: EPI5744602 |
| A/deshi-chicken/Bangladesh/AV-22-079931/2021 | HA: EPI5744605 / NA: EPI5744604 |
| A/deshi-chicken/Bangladesh/AV-22-079935/2021 | HA: EPI5744607 / NA: EPI5744606 |
| A/deshi-chicken/Bangladesh/AV-22-079938/2021 | HA: EPI5744609 / NA: EPI5744608 |
| A/deshi-chicken/Bangladesh/AV-22-079939/2021 | HA: EPI5744611 / NA: EPI5744610 |
| A/deshi-chicken/Bangladesh/AV-22-079943/2021 | HA: EPI5744613 / NA: EPI5744612 |
| A/deshi-chicken/Bangladesh/AV-22-079952/2021 | HA: EPI5744615 / NA: EPI5744614 |
| A/deshi-chicken/Bangladesh/AV-22-079954/2021 | HA: EPI5744617 / NA: EPI5744616 |
| A/deshi-chicken/Bangladesh/AV-22-079956/2021 | HA: EPI5744619 / NA: EPI5744618 |
| A/deshi-chicken/Bangladesh/AV-22-079971/2021 | HA: EPI5744621 / NA: EPI5744620 |
| A/sonali-chicken/Bangladesh/AV-22-005810/2021 | HA: EPI5744623 / NA: EPI5744622 |
| A/sonali-chicken/Bangladesh/AV-22-005817/2021 | HA: EPI5744625 / NA: EPI5744624 |
| A/sonali-chicken/Bangladesh/AV-22-006211/2021 | HA: EPI5744642 |
| A/sonali-chicken/Bangladesh/AV-22-006228/2021 | HA: EPI5744627 / NA: EPI5744626 |
| A/sonali-chicken/Bangladesh/AV-22-006230/2021 | HA: EPI5744629 / NA: EPI5744628 |
| A/sonali-chicken/Bangladesh/AV-22-006232/2021 | HA: EPI5744643 |
| A/sonali-chicken/Bangladesh/AV-22-006265/2021 | HA: EPI5744631 / NA: EPI5744630 |
| A/sonali-chicken/Bangladesh/AV-22-006269/2021 | HA: EPI5744633 / NA: EPI5744632 |
| A/sonali-chicken/Bangladesh/AV-22-006272/2021 | HA: EPI5744635 / NA: EPI5744634 |
| A/sonali-chicken/Bangladesh/AV-22-006298/2021 | HA: EPI5744637 / NA: EPI5744636 |
| A/sonali-chicken/Bangladesh/AV-22-006299/2021 | HA: EPI5744639 / NA: EPI5744638 |
| A/sonali-chicken/Bangladesh/AV-22-006305/2021 | HA: EPI5744641 / NA: EPI5744640 |
| A/sonali-chicken/Bangladesh/AV-22-006309/2021 | HA: EPI5744644 |
| A/sonali-chicken/Bangladesh/AV-22-006327/2021 | HA: EPI5744646 / NA: EPI5744645 |
| A/sonali-chicken/Bangladesh/AV-22-006331/2021 | HA: EPI5744648 / NA: EPI5744647 |
| A/sonali-chicken/Bangladesh/AV-22-006332/2021 | HA: EPI5744650 / NA: EPI5744649 |
| A/sonali-chicken/Bangladesh/AV-22-006344/2021 | HA: EPI5744652 / NA: EPI5744651 |
| A/sonali-chicken/Bangladesh/AV-22-006345/2021 | HA: EPI5744654 / NA: EPI5744653 |
| A/sonali-chicken/Bangladesh/AV-22-006354/2021 | HA: EPI5744656 / NA: EPI5744655 |
| A/sonali-chicken/Bangladesh/AV-22-006421/2021 | HA: EPI5744658 / NA: EPI5744657 |
| A/sonali-chicken/Bangladesh/AV-22-006424/2021 | HA: EPI5744660 / NA: EPI5744659 |
| A/sonali-chicken/Bangladesh/AV-22-006469/2021 | HA: EPI5744662 / NA: EPI5744661 |
| A/sonali-chicken/Bangladesh/AV-22-006470/2021 | HA: EPI5744664 / NA: EPI5744663 |
| A/sonali-chicken/Bangladesh/AV-22-006471/2021 | HA: EPI5744666 / NA: EPI5744665 |
| A/sonali-chicken/Bangladesh/AV-22-006472/2021 | HA: EPI5744668 / NA: EPI5744667 |
| A/sonali-chicken/Bangladesh/AV-22-006481/2021 | HA: EPI5744670 / NA: EPI5744669 |
| A/sonali-chicken/Bangladesh/AV-22-006503/2021 | HA: EPI5744672 / NA: EPI5744671 |
| A/sonali-chicken/Bangladesh/AV-22-006505/2021 | HA: EPI5744674 / NA: EPI5744673 |
| A/sonali-chicken/Bangladesh/AV-22-006507/2021 | HA: EPI5744676 / NA: EPI5744675 |
| A/sonali-chicken/Bangladesh/AV-22-006551/2021 | HA: EPI5744678 / NA: EPI5744677 |
| A/sonali-chicken/Bangladesh/AV-22-006561/2021 | HA: EPI5744680 / NA: EPI5744679 |
| A/sonali-chicken/Bangladesh/AV-22-006562/2021 | HA: EPI5744682 / NA: EPI5744681 |
| A/sonali-chicken/Bangladesh/AV-22-006585/2021 | HA: EPI5744684 / NA: EPI5744683 |
| A/sonali-chicken/Bangladesh/AV-22-006586/2021 | HA: EPI5744686 / NA: EPI5744685 |
| A/sonali-chicken/Bangladesh/AV-22-006619/2021 | HA: EPI5744688 / NA: EPI5744687 |
| A/sonali-chicken/Bangladesh/AV-22-006636/2021 | HA: EPI5744690 / NA: EPI5744689 |
| A/sonali-chicken/Bangladesh/AV-22-006658/2021 | HA: EPI5744692 / NA: EPI5744691 |
| A/sonali-chicken/Bangladesh/AV-22-006684/2021 | HA: EPI5744694 / NA: EPI5744693 |
| A/sonali-chicken/Bangladesh/AV-22-006685/2021 | HA: EPI5744696 / NA: EPI5744695 |
| A/sonali-chicken/Bangladesh/AV-22-079747/2021 | HA: EPI5744698 / NA: EPI5744697 |
| A/sonali-chicken/Bangladesh/AV-22-079748/2021 | HA: EPI5744700 / NA: EPI5744699 |
| A/sonali-chicken/Bangladesh/AV-22-079750/2021 | HA: EPI5744702 / NA: EPI5744701 |
| A/sonali-chicken/Bangladesh/AV-22-079752/2021 | HA: EPI5744704 / NA: EPI5744703 |
| A/sonali-chicken/Bangladesh/AV-22-079757/2021 | HA: EPI5744706 / NA: EPI5744705 |
| A/sonali-chicken/Bangladesh/AV-22-079758/2021 | HA: EPI5744708 / NA: EPI5744707 |
| A/sonali-chicken/Bangladesh/AV-22-079769/2021 | HA: EPI5744710 / NA: EPI5744709 |
| A/sonali-chicken/Bangladesh/AV-22-079770/2021 | HA: EPI5744712 / NA: EPI5744711 |
| A/sonali-chicken/Bangladesh/AV-22-079771/2021 | HA: EPI5744714 / NA: EPI5744713 |
| A/sonali-chicken/Bangladesh/AV-22-079774/2021 | HA: EPI5744716 / NA: EPI5744715 |
| A/sonali-chicken/Bangladesh/AV-22-079775/2021 | HA: EPI5744718 / NA: EPI5744717 |
| A/sonali-chicken/Bangladesh/AV-22-079776/2021 | HA: EPI5744720 / NA: EPI5744719 |
| A/sonali-chicken/Bangladesh/AV-22-079778/2021 | HA: EPI5744722 / NA: EPI5744721 |
| A/sonali-chicken/Bangladesh/AV-22-079783/2021 | HA: EPI5744724 / NA: EPI5744723 |
| A/sonali-chicken/Bangladesh/AV-22-079784/2021 | HA: EPI5744726 / NA: EPI5744725 |
| A/sonali-chicken/Bangladesh/AV-22-079785/2021 | HA: EPI5744728 / NA: EPI5744727 |
| A/sonali-chicken/Bangladesh/AV-22-079786/2021 | HA: EPI5744730 / NA: EPI5744729 |
| A/sonali-chicken/Bangladesh/AV-22-079790/2021 | HA: EPI5744732 / NA: EPI5744731 |
| A/sonali-chicken/Bangladesh/AV-22-079791/2021 | HA: EPI5744734 / NA: EPI5744733 |
| A/sonali-chicken/Bangladesh/AV-22-079792/2021 | HA: EPI5744736 / NA: EPI5744735 |
| A/sonali-chicken/Bangladesh/AV-22-079795/2021 | HA: EPI5744738 / NA: EPI5744737 |
| A/sonali-chicken/Bangladesh/AV-22-079797/2021 | HA: EPI5744740 / NA: EPI5744739 |
| A/sonali-chicken/Bangladesh/AV-22-079800/2021 | HA: EPI5744742 / NA: EPI5744741 |
| A/sonali-chicken/Bangladesh/AV-22-079801/2021 | HA: EPI5744744 / NA: EPI5744743 |
| A/sonali-chicken/Bangladesh/AV-22-079807/2021 | HA: EPI5744746 / NA: EPI5744745 |
| A/sonali-chicken/Bangladesh/AV-22-079812/2021 | HA: EPI5744748 / NA: EPI5744747 |
| A/sonali-chicken/Bangladesh/AV-22-079816/2021 | HA: EPI5744750 / NA: EPI5744749 |
| A/sonali-chicken/Bangladesh/AV-22-079817/2021 | HA: EPI5744752 / NA: EPI5744751 |
| A/sonali-chicken/Bangladesh/AV-22-079835/2021 | HA: EPI5744754 / NA: EPI5744753 |
| A/sonali-chicken/Bangladesh/AV-22-079838/2021 | HA: EPI5744756 / NA: EPI5744755 |
| A/sonali-chicken/Bangladesh/AV-22-079839/2021 | HA: EPI5744758 / NA: EPI5744757 |
| A/sonali-chicken/Bangladesh/AV-22-079856/2021 | HA: EPI5744760 / NA: EPI5744759 |
| A/sonali-chicken/Bangladesh/AV-22-079859/2021 | HA: EPI5744762 / NA: EPI5744761 |
| A/sonali-chicken/Bangladesh/AV-22-079861/2021 | HA: EPI5744764 / NA: EPI5744763 |
| A/sonali-chicken/Bangladesh/AV-22-079863/2021 | HA: EPI5744766 / NA: EPI5744765 |
| A/sonali-chicken/Bangladesh/AV-22-079877/2021 | HA: EPI5744768 / NA: EPI5744767 |
| A/sonali-chicken/Bangladesh/AV-22-079880/2021 | HA: EPI5744770 / NA: EPI5744769 |
| A/sonali-chicken/Bangladesh/AV-22-079881/2021 | HA: EPI5744772 / NA: EPI5744771 |
| A/sonali-chicken/Bangladesh/AV-22-079883/2021 | HA: EPI5744774 / NA: EPI5744773 |
| A/sonali-chicken/Bangladesh/AV-22-079884/2021 | HA: EPI5744776 / NA: EPI5744775 |
| A/sonali-chicken/Bangladesh/AV-22-079886/2021 | HA: EPI5744778 / NA: EPI5744777 |
| A/sonali-chicken/Bangladesh/AV-22-079895/2021 | HA: EPI5744780 / NA: EPI5744779 |
| A/sonali-chicken/Bangladesh/AV-22-079897/2021 | HA: EPI5744782 / NA: EPI5744781 |
| A/sonali-chicken/Bangladesh/AV-22-079921/2021 | HA: EPI5744784 / NA: EPI5744783 |
| A/sonali-chicken/Bangladesh/AV-22-079923/2021 | HA: EPI5744786 / NA: EPI5744785 |
| A/sonali-chicken/Bangladesh/AV-22-079924/2021 | HA: EPI5744788 / NA: EPI5744787 |
| A/sonali-chicken/Bangladesh/AV-22-079948/2021 | HA: EPI5744790 / NA: EPI5744789 |
| A/sonali-chicken/Bangladesh/AV-22-079949/2021 | HA: EPI5744792 / NA: EPI5744791 |
| A/sonali-chicken/Bangladesh/AV-22-079950/2021 | HA: EPI5744794 / NA: EPI5744793 |
| A/sonali-chicken/Bangladesh/AV-22-079951/2021 | HA: EPI5744796 / NA: EPI5744795 |
| A/sonali-chicken/Bangladesh/AV-22-079963/2021 | HA: EPI5744798 / NA: EPI5744797 |

**Table S2:** Quantification of HA-NA evolutionary discordance and reassortment status across Bangladeshi H9N2 poultry isolates. Viruses were ranked according to their HA-NA genomic discordance. Genetic mismatch score (0-1 composite score of spatial and distance discordance); procrustes residual (MDS alignment displacement); mean patristic discordance (average pairwise HA vs. NA branch-length variance); Spearman Rho (pairwise distance rank correlation); and local neighbour overlap (proportion of shared k=10 nearest patristic neighbours). Status reflects automated classification into true reassortant (mismatch score > Mean + 1.5 standard deviation (StD) and neighbour overlap ≤ 30%), co-evolved/variant neighbour overlap ≥ 60%), or intermediate/drift (neighbour overlap 31-59%).

| Rank | Strain | Genetic Mismatch Score | Procrustes Residual | Mean Patristic Discordance | Spearman Rho (ρ) | Local Neighbour Overlap | Status |
|---|---|---|---|---|---|---|---|
| 1 | A/sonali-chicken/Bangladesh/AV-22-079835/2021 | 0.9149 | 0.5694 | 0.3636 | 0.3456 | 0.3 | True Reassortant |
| 2 | A/sonali-chicken/Bangladesh/AV-22-079877/2021 | 0.889 | 0.552 | 0.3574 | 0.3245 | 0.1 | True Reassortant |
| 3 | A/broiler-chicken/Bangladesh/AV-22-006633/2021 | 0.8871 | 0.5834 | 0.3407 | 0.3236 | 0.1 | True Reassortant |
| 4 | A/deshi-chicken/Bangladesh/AV-22-006624/2021 | 0.87 | 0.5487 | 0.3482 | 0.337 | 0.2 | True Reassortant |
| 5 | A/broiler-chicken/Bangladesh/AV-22-006690/2021 | 0.8688 | 0.5485 | 0.3476 | 0.3352 | 0.2 | True Reassortant |
| 6 | A/broiler-chicken/Bangladesh/AV-22-006629/2021 | 0.8686 | 0.5485 | 0.3475 | 0.3353 | 0.2 | True Reassortant |
| 7 | A/sonali-chicken/Bangladesh/AV-22-079881/2021 | 0.845 | 0.5547 | 0.3309 | 0.3078 | 0.3 | True Reassortant |
| 8 | A/sonali-chicken/Bangladesh/AV-22-079883/2021 | 0.8445 | 0.5547 | 0.3306 | 0.3078 | 0.3 | True Reassortant |
| 9 | A/broiler-chicken/Bangladesh/AV-22-006680/2021 | 0.8332 | 0.551 | 0.326 | 0.3375 | 0.1 | True Reassortant |
| 10 | A/sonali-chicken/Bangladesh/AV-22-006619/2021 | 0.8327 | 0.551 | 0.3257 | 0.3375 | 0.2 | True Reassortant |
| 11 | A/sonali-chicken/Bangladesh/AV-22-079884/2021 | 0.8289 | 0.5275 | 0.3352 | 0.3219 | 0.1 | True Reassortant |
| 12 | A/broiler-chicken/Bangladesh/AV-22-079869/2021 | 0.8267 | 0.529 | 0.3332 | 0.3335 | 0.2 | True Reassortant |
| 13 | A/broiler-chicken/Bangladesh/AV-22-079867/2021 | 0.8261 | 0.5229 | 0.3359 | 0.3336 | 0 | True Reassortant |
| 14 | A/broiler-chicken/Bangladesh/AV-22-079875/2021 | 0.8077 | 0.5514 | 0.3112 | 0.3078 | 0.3 | True Reassortant |
| 15 | A/sonali-chicken/Bangladesh/AV-22-079880/2021 | 0.8077 | 0.5514 | 0.3112 | 0.3078 | 0.4 | Intermediate / Drift |
| 16 | A/broiler-chicken/Bangladesh/AV-22-079728/2021 | 0.8016 | 0.4706 | 0.3479 | -0.1528 | 0.1 | True Reassortant |
| 17 | A/sonali-chicken/Bangladesh/AV-22-006481/2021 | 0.7996 | 0.4685 | 0.3478 | -0.152 | 0.1 | True Reassortant |
| 18 | A/deshi-chicken/Bangladesh/AV-22-079971/2021 | 0.7844 | 0.6049 | 0.2713 | 0.0144 | 0 | True Reassortant |
| 19 | A/broiler-chicken/Bangladesh/AV-22-006674/2021 | 0.746 | 0.5894 | 0.257 | 0.2806 | 0.1 | True Reassortant |
| 20 | A/sonali-chicken/Bangladesh/AV-22-006685/2021 | 0.7458 | 0.5036 | 0.2996 | 0.3221 | 0.6 | Co-Evolved / Variant |
| 21 | A/sonali-chicken/Bangladesh/AV-22-079863/2021 | 0.7419 | 0.5347 | 0.2819 | 0.3175 | 0 | True Reassortant |
| 22 | A/broiler-chicken/Bangladesh/AV-22-006196/2021 | 0.7401 | 0.3395 | 0.3779 | -0.0817 | 0.1 | True Reassortant |
| 23 | A/broiler-chicken/Bangladesh/AV-22-079850/2021 | 0.7391 | 0.6163 | 0.2397 | 0.3066 | 0.1 | Intermediate / Drift |
| 24 | A/sonali-chicken/Bangladesh/AV-22-079783/2021 | 0.7317 | 0.3276 | 0.379 | -0.0876 | 0.2 | Intermediate / Drift |
| 25 | A/broiler-chicken/Bangladesh/AV-22-079870/2021 | 0.731 | 0.4875 | 0.2991 | 0.3366 | 0 | Intermediate / Drift |
| 26 | A/broiler-chicken/Bangladesh/AV-22-005799/2021 | 0.7294 | 0.3095 | 0.3867 | -0.0973 | 0.2 | Intermediate / Drift |
| 27 | A/broiler-chicken/Bangladesh/AV-22-006689/2021 | 0.717 | 0.4945 | 0.2876 | 0.3193 | 0.4 | Intermediate / Drift |
| 28 | A/deshi-chicken/Bangladesh/AV-22-079938/2021 | 0.7144 | 0.5448 | 0.2611 | 0.1931 | 0.1 | Intermediate / Drift |
| 29 | A/deshi-chicken/Bangladesh/AV-22-079943/2021 | 0.7144 | 0.5448 | 0.2611 | 0.1937 | 0.1 | Intermediate / Drift |
| 30 | A/broiler-chicken/Bangladesh/AV-22-006654/2021 | 0.7133 | 0.478 | 0.2937 | 0.3107 | 0.3 | Intermediate / Drift |
| 31 | A/sonali-chicken/Bangladesh/AV-22-079839/2021 | 0.7119 | 0.6208 | 0.2219 | 0.3066 | 0.1 | Intermediate / Drift |
| 32 | A/broiler-chicken/Bangladesh/AV-22-006614/2021 | 0.7036 | 0.5383 | 0.2582 | 0.2964 | 0.1 | Intermediate / Drift |
| 33 | A/deshi-chicken/Bangladesh/AV-22-006213/2021 | 0.6957 | 0.2749 | 0.3846 | -0.0969 | 0.2 | Intermediate / Drift |
| 34 | A/sonali-chicken/Bangladesh/AV-22-079861/2021 | 0.6928 | 0.591 | 0.2258 | 0.3073 | 0.2 | Intermediate / Drift |
| 35 | A/broiler-chicken/Bangladesh/AV-22-079854/2021 | 0.6928 | 0.591 | 0.2258 | 0.3074 | 0.2 | Intermediate / Drift |
| 36 | A/broiler-chicken/Bangladesh/AV-22-079852/2021 | 0.6904 | 0.5895 | 0.2252 | 0.3055 | 0.1 | Intermediate / Drift |
| 37 | A/broiler-chicken/Bangladesh/AV-22-079821/2021 | 0.6852 | 0.4739 | 0.2797 | 0.3097 | 0.2 | Intermediate / Drift |
| 38 | A/broiler-chicken/Bangladesh/AV-22-079823/2021 | 0.6852 | 0.4739 | 0.2797 | 0.3092 | 0.2 | Intermediate / Drift |
| 39 | A/broiler-chicken/Bangladesh/AV-22-079824/2021 | 0.6852 | 0.4739 | 0.2797 | 0.3088 | 0.3 | Intermediate / Drift |
| 40 | A/broiler-chicken/Bangladesh/AV-22-006374/2021 | 0.6814 | 0.5847 | 0.2224 | 0.302 | 0.1 | Intermediate / Drift |
| 41 | A/broiler-chicken/Bangladesh/AV-22-006682/2021 | 0.6789 | 0.4823 | 0.2719 | 0.3186 | 0.4 | Intermediate / Drift |
| 42 | A/sonali-chicken/Bangladesh/AV-22-006684/2021 | 0.6789 | 0.4823 | 0.2719 | 0.3153 | 0.2 | Intermediate / Drift |
| 43 | A/sonali-chicken/Bangladesh/AV-22-079856/2021 | 0.6777 | 0.4731 | 0.2758 | 0.3048 | 0 | Intermediate / Drift |
| 44 | A/sonali-chicken/Bangladesh/AV-22-079859/2021 | 0.6777 | 0.4731 | 0.2758 | 0.3041 | 0 | Intermediate / Drift |
| 45 | A/sonali-chicken/Bangladesh/AV-22-006658/2021 | 0.6554 | 0.4796 | 0.2598 | 0.3192 | 0.4 | Intermediate / Drift |
| 46 | A/sonali-chicken/Bangladesh/AV-22-079838/2021 | 0.6497 | 0.4807 | 0.256 | 0.3137 | 0.1 | Intermediate / Drift |
| 47 | A/broiler-chicken/Bangladesh/AV-22-006662/2021 | 0.6479 | 0.48 | 0.2553 | 0.3187 | 0.5 | Intermediate / Drift |
| 48 | A/sonali-chicken/Bangladesh/AV-22-079886/2021 | 0.6436 | 0.5634 | 0.2114 | 0.3037 | 0.2 | Intermediate / Drift |
| 49 | A/broiler-chicken/Bangladesh/AV-22-079868/2021 | 0.6435 | 0.5633 | 0.2114 | 0.3035 | 0.1 | Intermediate / Drift |
| 50 | A/broiler-chicken/Bangladesh/AV-22-079904/2021 | 0.6356 | 0.2305 | 0.3723 | -0.3216 | 0.1 | Intermediate / Drift |
| 51 | A/broiler-chicken/Bangladesh/AV-22-079833/2021 | 0.6319 | 0.5422 | 0.2152 | 0.2943 | 0.1 | Intermediate / Drift |
| 52 | A/deshi-chicken/Bangladesh/AV-22-006383/2021 | 0.5967 | 0.4306 | 0.2506 | -0.1518 | 0 | Intermediate / Drift |
| 53 | A/broiler-chicken/Bangladesh/AV-22-006392/2021 | 0.5906 | 0.4342 | 0.2453 | -0.1519 | 0 | Intermediate / Drift |
| 54 | A/deshi-chicken/Bangladesh/AV-22-079939/2021 | 0.5742 | 0.4644 | 0.2209 | -0.1976 | 0 | Intermediate / Drift |
| 55 | A/sonali-chicken/Bangladesh/AV-22-079950/2021 | 0.5666 | 0.4177 | 0.2398 | -0.1929 | 0.1 | Intermediate / Drift |
| 56 | A/deshi-chicken/Bangladesh/AV-22-079743/2021 | 0.5447 | 0.1016 | 0.3844 | -0.0628 | 0.1 | Intermediate / Drift |
| 57 | A/broiler-chicken/Bangladesh/AV-22-079906/2021 | 0.5401 | 0.4602 | 0.2035 | -0.0999 | 0 | Intermediate / Drift |
| 58 | A/sonali-chicken/Bangladesh/AV-22-079963/2021 | 0.5377 | 0.4061 | 0.229 | -0.0562 | 0.2 | Intermediate / Drift |
| 59 | A/sonali-chicken/Bangladesh/AV-22-006505/2021 | 0.537 | 0.3124 | 0.2752 | -0.0817 | 0.1 | Intermediate / Drift |
| 60 | A/deshi-chicken/Bangladesh/AV-22-006410/2021 | 0.5358 | 0.1302 | 0.3651 | -0.07 | 0.2 | Intermediate / Drift |
| 61 | A/broiler-chicken/Bangladesh/AV-22-006400/2021 | 0.5358 | 0.1302 | 0.3651 | -0.07 | 0.2 | Intermediate / Drift |
| 62 | A/sonali-chicken/Bangladesh/AV-22-079949/2021 | 0.5335 | 0.4172 | 0.2211 | -0.0536 | 0.3 | Intermediate / Drift |
| 63 | A/deshi-chicken/Bangladesh/AV-22-079952/2021 | 0.5333 | 0.359 | 0.2499 | -0.1937 | 0.1 | Intermediate / Drift |
| 64 | A/broiler-chicken/Bangladesh/AV-22-006661/2021 | 0.5321 | 0.3983 | 0.2297 | -0.2011 | 0 | Intermediate / Drift |
| 65 | A/broiler-chicken/Bangladesh/AV-22-006293/2021 | 0.5222 | 0.3931 | 0.2266 | -0.0573 | 0.1 | Intermediate / Drift |
| 66 | A/sonali-chicken/Bangladesh/AV-22-006228/2021 | 0.5214 | 0.093 | 0.3754 | -0.0787 | 0.2 | Intermediate / Drift |
| 67 | A/deshi-chicken/Bangladesh/AV-22-006612/2021 | 0.5159 | 0.3406 | 0.2491 | -0.1619 | 0 | Intermediate / Drift |
| 68 | A/deshi-chicken/Bangladesh/AV-22-006449/2021 | 0.5091 | 0.1158 | 0.357 | -0.0785 | 0.2 | Intermediate / Drift |
| 69 | A/deshi-chicken/Bangladesh/AV-22-079954/2021 | 0.5091 | 0.4088 | 0.2113 | -0.0534 | 0.3 | Intermediate / Drift |
| 70 | A/broiler-chicken/Bangladesh/AV-22-079772/2021 | 0.507 | 0.2105 | 0.3087 | -0.2842 | 0.1 | Intermediate / Drift |
| 71 | A/sonali-chicken/Bangladesh/AV-22-079948/2021 | 0.4977 | 0.406 | 0.2062 | -0.0535 | 0.3 | Intermediate / Drift |
| 72 | A/deshi-chicken/Bangladesh/AV-22-006523/2021 | 0.4968 | 0.3935 | 0.2119 | -0.088 | 0.1 | Intermediate / Drift |
| 73 | A/sonali-chicken/Bangladesh/AV-22-006327/2021 | 0.4944 | 0.3889 | 0.2128 | -0.0516 | 0.2 | Intermediate / Drift |
| 74 | A/sonali-chicken/Bangladesh/AV-22-006331/2021 | 0.4944 | 0.3889 | 0.2128 | -0.0516 | 0.2 | Intermediate / Drift |
| 75 | A/deshi-chicken/Bangladesh/AV-22-006338/2021 | 0.4938 | 0.3882 | 0.2128 | -0.0586 | 0 | Intermediate / Drift |
| 76 | A/sonali-chicken/Bangladesh/AV-22-006332/2021 | 0.4938 | 0.3882 | 0.2128 | -0.0569 | 0.1 | Intermediate / Drift |
| 77 | A/deshi-chicken/Bangladesh/AV-22-006408/2021 | 0.4905 | 0.3822 | 0.2139 | -0.154 | 0 | Intermediate / Drift |
| 78 | A/broiler-chicken/Bangladesh/AV-22-006342/2021 | 0.4873 | 0.4008 | 0.2028 | -0.0879 | 0.1 | Intermediate / Drift |
| 79 | A/broiler-chicken/Bangladesh/AV-22-006531/2021 | 0.4758 | 0.1648 | 0.3136 | -0.28 | 0.1 | Intermediate / Drift |
| 80 | A/sonali-chicken/Bangladesh/AV-22-079786/2021 | 0.4714 | 0.3614 | 0.2133 | -0.1158 | 0 | Intermediate / Drift |
| 81 | A/broiler-chicken/Bangladesh/AV-22-079941/2021 | 0.4635 | 0.3256 | 0.2266 | -0.0424 | 0.3 | Intermediate / Drift |
| 82 | A/deshi-chicken/Bangladesh/AV-22-079744/2021 | 0.4604 | 0.1052 | 0.3344 | -0.0628 | 0.1 | Intermediate / Drift |
| 83 | A/broiler-chicken/Bangladesh/AV-22-079942/2021 | 0.4576 | 0.3518 | 0.2102 | -0.0431 | 0.2 | Intermediate / Drift |
| 84 | A/broiler-chicken/Bangladesh/AV-22-079741/2021 | 0.4507 | 0.2586 | 0.2526 | -0.1904 | 0 | Intermediate / Drift |
| 85 | A/sonali-chicken/Bangladesh/AV-22-079747/2021 | 0.4493 | 0.2278 | 0.2671 | -0.2015 | 0.2 | Intermediate / Drift |
| 86 | A/sonali-chicken/Bangladesh/AV-22-079951/2021 | 0.4476 | 0.3453 | 0.2077 | -0.0448 | 0.3 | Intermediate / Drift |
| 87 | A/broiler-chicken/Bangladesh/AV-22-079940/2021 | 0.4469 | 0.3286 | 0.2156 | -0.0431 | 0.2 | Intermediate / Drift |
| 88 | A/sonali-chicken/Bangladesh/AV-22-079774/2021 | 0.4464 | 0.3299 | 0.2147 | -0.0731 | 0.3 | Intermediate / Drift |
| 89 | A/broiler-chicken/Bangladesh/AV-22-006379/2021 | 0.4415 | 0.3071 | 0.2232 | 0.0923 | 0.1 | Intermediate / Drift |
| 90 | A/deshi-chicken/Bangladesh/AV-22-079740/2021 | 0.4363 | 0.2374 | 0.2549 | -0.177 | 0 | Intermediate / Drift |
| 91 | A/sonali-chicken/Bangladesh/AV-22-079775/2021 | 0.4352 | 0.3099 | 0.2182 | -0.0731 | 0.3 | Intermediate / Drift |
| 92 | A/sonali-chicken/Bangladesh/AV-22-006354/2021 | 0.4316 | 0.2974 | 0.2224 | -0.19 | 0.1 | Intermediate / Drift |
| 93 | A/sonali-chicken/Bangladesh/AV-22-006345/2021 | 0.4285 | 0.2228 | 0.2577 | -0.2014 | 0.2 | Intermediate / Drift |
| 94 | A/sonali-chicken/Bangladesh/AV-22-079897/2021 | 0.4268 | 0.2784 | 0.2291 | -0.0731 | 0.3 | Intermediate / Drift |
| 95 | A/deshi-chicken/Bangladesh/AV-22-079742/2021 | 0.4247 | 0.1613 | 0.2861 | -0.2077 | 0 | Intermediate / Drift |
| 96 | A/sonali-chicken/Bangladesh/AV-22-079895/2021 | 0.4239 | 0.2782 | 0.2275 | -0.0731 | 0.2 | Intermediate / Drift |
| 97 | A/deshi-chicken/Bangladesh/AV-22-006317/2021 | 0.4226 | 0.2335 | 0.249 | 0.0984 | 0.3 | Intermediate / Drift |
| 98 | A/sonali-chicken/Bangladesh/AV-22-006562/2021 | 0.413 | 0.1814 | 0.2694 | -0.1899 | 0.1 | Intermediate / Drift |
| 99 | A/deshi-chicken/Bangladesh/AV-22-079931/2021 | 0.4125 | 0.3162 | 0.2021 | 0.0712 | 0 | Intermediate / Drift |
| 100 | A/deshi-chicken/Bangladesh/AV-22-079842/2021 | 0.4124 | 0.2891 | 0.2155 | -0.0242 | 0 | Intermediate / Drift |
| 101 | A/deshi-chicken/Bangladesh/AV-22-079844/2021 | 0.4121 | 0.3085 | 0.2057 | -0.0242 | 0.1 | Intermediate / Drift |
| 102 | A/sonali-chicken/Bangladesh/AV-22-079758/2021 | 0.4111 | 0.1935 | 0.2623 | -0.1322 | 0.3 | Intermediate / Drift |
| 103 | A/deshi-chicken/Bangladesh/AV-22-006357/2021 | 0.4079 | 0.2174 | 0.2486 | -0.2038 | 0.1 | Intermediate / Drift |
| 104 | A/deshi-chicken/Bangladesh/AV-22-006433/2021 | 0.4073 | 0.2091 | 0.2524 | -0.1772 | 0 | Intermediate / Drift |
| 105 | A/broiler-chicken/Bangladesh/AV-22-006323/2021 | 0.4073 | 0.231 | 0.2415 | -0.171 | 0.2 | Intermediate / Drift |
| 106 | A/deshi-chicken/Bangladesh/AV-22-079889/2021 | 0.4051 | 0.1735 | 0.2688 | -0.0797 | 0.1 | Intermediate / Drift |
| 107 | A/deshi-chicken/Bangladesh/AV-22-006487/2021 | 0.404 | 0.1727 | 0.2686 | -0.1604 | 0.1 | Intermediate / Drift |
| 108 | A/sonali-chicken/Bangladesh/AV-22-079807/2021 | 0.4036 | 0.2074 | 0.2511 | -0.171 | 0.2 | Intermediate / Drift |
| 109 | A/sonali-chicken/Bangladesh/AV-22-006298/2021 | 0.4028 | 0.3105 | 0.1994 | 0.1072 | 0.6 | Co-Evolved / Variant |
| 110 | A/deshi-chicken/Bangladesh/AV-22-006388/2021 | 0.4023 | 0.1603 | 0.2738 | -0.0615 | 0 | Intermediate / Drift |
| 111 | A/broiler-chicken/Bangladesh/AV-22-006607/2021 | 0.4022 | 0.2997 | 0.2044 | -0.0489 | 0.1 | Intermediate / Drift |
| 112 | A/deshi-chicken/Bangladesh/AV-22-079929/2021 | 0.3975 | 0.2929 | 0.2051 | 0.0711 | 0 | Intermediate / Drift |
| 113 | A/deshi-chicken/Bangladesh/AV-22-079843/2021 | 0.3966 | 0.2488 | 0.2265 | -0.1559 | 0.1 | Intermediate / Drift |
| 114 | A/broiler-chicken/Bangladesh/AV-22-006500/2021 | 0.3932 | 0.2423 | 0.2278 | -0.0841 | 0 | Intermediate / Drift |
| 115 | A/broiler-chicken/Bangladesh/AV-22-006579/2021 | 0.3929 | 0.2963 | 0.2008 | 0.1072 | 0.5 | Intermediate / Drift |
| 116 | A/sonali-chicken/Bangladesh/AV-22-006551/2021 | 0.3927 | 0.181 | 0.258 | -0.171 | 0.2 | Intermediate / Drift |
| 117 | A/broiler-chicken/Bangladesh/AV-22-006294/2021 | 0.3912 | 0.2676 | 0.2141 | 0.0645 | 0 | Intermediate / Drift |
| 118 | A/sonali-chicken/Bangladesh/AV-22-006561/2021 | 0.3909 | 0.1765 | 0.2592 | -0.1899 | 0 | Intermediate / Drift |
| 119 | A/sonali-chicken/Bangladesh/AV-22-079800/2021 | 0.3901 | 0.294 | 0.2003 | 0.1055 | 0.2 | Intermediate / Drift |
| 120 | A/deshi-chicken/Bangladesh/AV-22-006496/2021 | 0.39 | 0.1662 | 0.2638 | -0.1917 | 0 | Intermediate / Drift |
| 121 | A/sonali-chicken/Bangladesh/AV-22-006305/2021 | 0.3887 | 0.2215 | 0.2356 | 0.0997 | 0.3 | Intermediate / Drift |
| 122 | A/sonali-chicken/Bangladesh/AV-22-079752/2021 | 0.3883 | 0.1645 | 0.2637 | -0.1322 | 0.3 | Intermediate / Drift |
| 123 | A/sonali-chicken/Bangladesh/AV-22-079750/2021 | 0.3872 | 0.1887 | 0.251 | -0.1333 | 0.2 | Intermediate / Drift |
| 124 | A/sonali-chicken/Bangladesh/AV-22-006471/2021 | 0.3866 | 0.2822 | 0.2042 | 0.1049 | 0.2 | Intermediate / Drift |
| 125 | A/deshi-chicken/Bangladesh/AV-22-006340/2021 | 0.3857 | 0.2816 | 0.204 | 0.1047 | 0.3 | Intermediate / Drift |
| 126 | A/broiler-chicken/Bangladesh/AV-22-006556/2021 | 0.3857 | 0.2816 | 0.204 | 0.1047 | 0.3 | Intermediate / Drift |
| 127 | A/sonali-chicken/Bangladesh/AV-22-079801/2021 | 0.3851 | 0.2895 | 0.1997 | 0.1067 | 0.3 | Intermediate / Drift |
| 128 | A/deshi-chicken/Bangladesh/AV-22-006596/2021 | 0.3847 | 0.2728 | 0.2078 | 0.0869 | 0 | Intermediate / Drift |
| 129 | A/deshi-chicken/Bangladesh/AV-22-006597/2021 | 0.3831 | 0.2534 | 0.2165 | 0.1071 | 0.4 | Intermediate / Drift |
| 130 | A/sonali-chicken/Bangladesh/AV-22-079785/2021 | 0.3777 | 0.2028 | 0.2386 | -0.1678 | 0.2 | Intermediate / Drift |
| 131 | A/sonali-chicken/Bangladesh/AV-22-006470/2021 | 0.3776 | 0.1319 | 0.2738 | -0.169 | 0.4 | Intermediate / Drift |
| 132 | A/deshi-chicken/Bangladesh/AV-22-079935/2021 | 0.3759 | 0.2576 | 0.2103 | 0.1049 | 0.2 | Intermediate / Drift |
| 133 | A/broiler-chicken/Bangladesh/AV-22-006455/2021 | 0.3757 | 0.2576 | 0.2102 | 0.1018 | 0.1 | Intermediate / Drift |
| 134 | A/broiler-chicken/Bangladesh/AV-22-006459/2021 | 0.3757 | 0.2576 | 0.2102 | 0.1011 | 0.2 | Intermediate / Drift |
| 135 | A/sonali-chicken/Bangladesh/AV-22-079797/2021 | 0.3725 | 0.2624 | 0.206 | 0.1079 | 0.4 | Intermediate / Drift |
| 136 | A/sonali-chicken/Bangladesh/AV-22-006586/2021 | 0.3701 | 0.2714 | 0.2001 | 0.1074 | 0.4 | Intermediate / Drift |
| 137 | A/broiler-chicken/Bangladesh/AV-22-006570/2021 | 0.3699 | 0.2714 | 0.2 | 0.1028 | 0.2 | Intermediate / Drift |
| 138 | A/sonali-chicken/Bangladesh/AV-22-006585/2021 | 0.3699 | 0.2714 | 0.2 | 0.1064 | 0.4 | Intermediate / Drift |
| 139 | A/sonali-chicken/Bangladesh/AV-22-006299/2021 | 0.3699 | 0.2714 | 0.2 | 0.1074 | 0.5 | Intermediate / Drift |
| 140 | A/deshi-chicken/Bangladesh/AV-22-006442/2021 | 0.3681 | 0.1947 | 0.2371 | -0.326 | 0 | Intermediate / Drift |
| 141 | A/sonali-chicken/Bangladesh/AV-22-079778/2021 | 0.3521 | 0.1934 | 0.2286 | -0.074 | 0.2 | Intermediate / Drift |
| 142 | A/deshi-chicken/Bangladesh/AV-22-079788/2021 | 0.3476 | 0.1732 | 0.2361 | -0.1767 | 0 | Intermediate / Drift |
| 143 | A/deshi-chicken/Bangladesh/AV-22-079830/2021 | 0.3463 | 0.1767 | 0.2336 | -0.0798 | 0 | Intermediate / Drift |
| 144 | A/sonali-chicken/Bangladesh/AV-22-079776/2021 | 0.3411 | 0.1645 | 0.2367 | -0.074 | 0.1 | Intermediate / Drift |
| 145 | A/sonali-chicken/Bangladesh/AV-22-006424/2021 | 0.339 | 0.2139 | 0.2109 | -0.15 | 0 | Intermediate / Drift |
| 146 | A/deshi-chicken/Bangladesh/AV-22-079781/2021 | 0.3357 | 0.1231 | 0.2542 | -0.1598 | 0.2 | Intermediate / Drift |
| 147 | A/deshi-chicken/Bangladesh/AV-22-079779/2021 | 0.3357 | 0.1231 | 0.2542 | -0.1598 | 0.2 | Intermediate / Drift |
| 148 | A/broiler-chicken/Bangladesh/AV-22-005796/2021 | 0.3351 | 0.0484 | 0.291 | 0.0969 | 0.2 | Intermediate / Drift |
| 149 | A/sonali-chicken/Bangladesh/AV-22-005817/2021 | 0.3351 | 0.0484 | 0.291 | 0.097 | 0.2 | Intermediate / Drift |
| 150 | A/sonali-chicken/Bangladesh/AV-22-005810/2021 | 0.3351 | 0.0484 | 0.291 | 0.0967 | 0.3 | Intermediate / Drift |
| 151 | A/broiler-chicken/Bangladesh/AV-22-079919/2021 | 0.3349 | 0.0685 | 0.2809 | -0.1036 | 0.1 | Intermediate / Drift |
| 152 | A/deshi-chicken/Bangladesh/AV-22-006494/2021 | 0.3332 | 0.1379 | 0.2454 | -0.169 | 0.3 | Intermediate / Drift |
| 153 | A/broiler-chicken/Bangladesh/AV-22-079959/2021 | 0.333 | 0.1529 | 0.2378 | -0.174 | 0 | Intermediate / Drift |
| 154 | A/deshi-chicken/Bangladesh/AV-22-006289/2021 | 0.3286 | 0.1244 | 0.2495 | -0.1727 | 0.2 | Intermediate / Drift |
| 155 | A/deshi-chicken/Bangladesh/AV-22-006287/2021 | 0.3286 | 0.1244 | 0.2495 | -0.1727 | 0.2 | Intermediate / Drift |
| 156 | A/broiler-chicken/Bangladesh/AV-22-079936/2021 | 0.3264 | 0.1118 | 0.2545 | -0.0866 | 0 | Intermediate / Drift |
| 157 | A/broiler-chicken/Bangladesh/AV-22-006463/2021 | 0.3223 | 0.1055 | 0.2553 | -0.0796 | 0.1 | Intermediate / Drift |
| 158 | A/deshi-chicken/Bangladesh/AV-22-079780/2021 | 0.3201 | 0.0898 | 0.2618 | -0.082 | 0.1 | Intermediate / Drift |
| 159 | A/broiler-chicken/Bangladesh/AV-22-006378/2021 | 0.3194 | 0.0959 | 0.2584 | -0.1686 | 0.1 | Intermediate / Drift |
| 160 | A/sonali-chicken/Bangladesh/AV-22-079791/2021 | 0.3129 | 0.2018 | 0.202 | 0.0432 | 0 | Intermediate / Drift |
| 161 | A/deshi-chicken/Bangladesh/AV-22-079890/2021 | 0.3114 | 0.0959 | 0.2538 | 0.0776 | 0 | Intermediate / Drift |
| 162 | A/broiler-chicken/Bangladesh/AV-22-006391/2021 | 0.3092 | 0.1676 | 0.2169 | -0.1267 | 0.1 | Intermediate / Drift |
| 163 | A/deshi-chicken/Bangladesh/AV-22-079903/2021 | 0.3065 | 0.1926 | 0.2029 | -0.0019 | 0.1 | Intermediate / Drift |
| 164 | A/deshi-chicken/Bangladesh/AV-22-079787/2021 | 0.3054 | 0.1699 | 0.2136 | 0.0379 | 0.3 | Intermediate / Drift |
| 165 | A/deshi-chicken/Bangladesh/AV-22-006492/2021 | 0.3033 | 0.1417 | 0.2264 | -0.169 | 0.3 | Intermediate / Drift |
| 166 | A/deshi-chicken/Bangladesh/AV-22-079737/2021 | 0.3002 | 0.0762 | 0.2572 | 0.0984 | 0.2 | Intermediate / Drift |
| 167 | A/sonali-chicken/Bangladesh/AV-22-006344/2021 | 0.2987 | 0.1716 | 0.2089 | 0.1403 | 0.2 | Intermediate / Drift |
| 168 | A/sonali-chicken/Bangladesh/AV-22-079792/2021 | 0.2927 | 0.1734 | 0.2046 | 0.0432 | 0.1 | Intermediate / Drift |
| 169 | A/sonali-chicken/Bangladesh/AV-22-006469/2021 | 0.292 | 0.1579 | 0.2119 | -0.169 | 0.3 | Intermediate / Drift |
| 170 | A/broiler-chicken/Bangladesh/AV-22-079729/2021 | 0.2916 | 0.0456 | 0.2675 | 0.0964 | 0.2 | Intermediate / Drift |
| 171 | A/broiler-chicken/Bangladesh/AV-22-079872/2021 | 0.29 | 0.1789 | 0.2003 | 0.0776 | 0 | Intermediate / Drift |
| 172 | A/sonali-chicken/Bangladesh/AV-22-006272/2021 | 0.2883 | 0.0659 | 0.2555 | 0.0963 | 0.3 | Intermediate / Drift |
| 173 | A/deshi-chicken/Bangladesh/AV-22-006359/2021 | 0.2846 | 0.1287 | 0.2222 | -0.0016 | 0.2 | Intermediate / Drift |
| 174 | A/sonali-chicken/Bangladesh/AV-22-006230/2021 | 0.2836 | 0.1597 | 0.2062 | -0.0015 | 0.2 | Intermediate / Drift |
| 175 | A/sonali-chicken/Bangladesh/AV-22-006503/2021 | 0.2828 | 0.1185 | 0.2262 | 0.0161 | 0 | Intermediate / Drift |
| 176 | A/sonali-chicken/Bangladesh/AV-22-079769/2021 | 0.2826 | 0.0602 | 0.2551 | 0.0772 | 0.2 | Intermediate / Drift |
| 177 | A/sonali-chicken/Bangladesh/AV-22-079771/2021 | 0.2814 | 0.0602 | 0.2544 | 0.0772 | 0.1 | Intermediate / Drift |
| 178 | A/sonali-chicken/Bangladesh/AV-22-079770/2021 | 0.2814 | 0.0602 | 0.2544 | 0.0772 | 0.1 | Intermediate / Drift |
| 179 | A/sonali-chicken/Bangladesh/AV-22-006421/2021 | 0.2782 | 0.0861 | 0.2397 | -0.131 | 0.1 | Intermediate / Drift |
| 180 | A/broiler-chicken/Bangladesh/AV-22-006194/2021 | 0.2777 | 0.0573 | 0.2537 | 0.0989 | 0.2 | Intermediate / Drift |
| 181 | A/deshi-chicken/Bangladesh/AV-22-079759/2021 | 0.2766 | 0.138 | 0.213 | 0.1143 | 0 | Intermediate / Drift |
| 182 | A/sonali-chicken/Bangladesh/AV-22-006507/2021 | 0.2735 | 0.0581 | 0.2509 | 0.1405 | 0.1 | Intermediate / Drift |
| 183 | A/broiler-chicken/Bangladesh/AV-22-079918/2021 | 0.2728 | 0.1465 | 0.2066 | 0.0301 | 0.4 | Intermediate / Drift |
| 184 | A/sonali-chicken/Bangladesh/AV-22-079748/2021 | 0.2701 | 0.0762 | 0.24 | 0.095 | 0 | Intermediate / Drift |
| 185 | A/broiler-chicken/Bangladesh/AV-22-079745/2021 | 0.2678 | 0.0962 | 0.2287 | 0.1196 | 0.1 | Intermediate / Drift |
| 186 | A/deshi-chicken/Bangladesh/AV-22-006247/2021 | 0.2657 | 0.1206 | 0.2154 | 0.0161 | 0.2 | Intermediate / Drift |
| 187 | A/sonali-chicken/Bangladesh/AV-22-079812/2021 | 0.2652 | 0.0981 | 0.2263 | 0.0983 | 0.2 | Intermediate / Drift |
| 188 | A/broiler-chicken/Bangladesh/AV-22-079957/2021 | 0.265 | 0.1624 | 0.1942 | -0.0355 | 0 | Intermediate / Drift |
| 189 | A/sonali-chicken/Bangladesh/AV-22-079924/2021 | 0.2632 | 0.0666 | 0.2408 | 0.012 | 0.3 | Intermediate / Drift |
| 190 | A/sonali-chicken/Bangladesh/AV-22-079923/2021 | 0.2632 | 0.0666 | 0.2408 | 0.0118 | 0.3 | Intermediate / Drift |
| 191 | A/sonali-chicken/Bangladesh/AV-22-079921/2021 | 0.2632 | 0.0666 | 0.2408 | 0.0118 | 0.3 | Intermediate / Drift |
| 192 | A/broiler-chicken/Bangladesh/AV-22-006219/2021 | 0.2621 | 0.0881 | 0.2295 | 0.0781 | 0.2 | Intermediate / Drift |
| 193 | A/deshi-chicken/Bangladesh/AV-22-006256/2021 | 0.2621 | 0.0881 | 0.2295 | 0.078 | 0.2 | Intermediate / Drift |
| 194 | A/broiler-chicken/Bangladesh/AV-22-006390/2021 | 0.2617 | 0.1236 | 0.2116 | 0.1196 | 0.1 | Intermediate / Drift |
| 195 | A/deshi-chicken/Bangladesh/AV-22-006407/2021 | 0.2605 | 0.0478 | 0.2486 | 0.0895 | 0.1 | Intermediate / Drift |
| 196 | A/sonali-chicken/Bangladesh/AV-22-079816/2021 | 0.2554 | 0.1343 | 0.2027 | 0.0984 | 0.2 | Intermediate / Drift |
| 197 | A/deshi-chicken/Bangladesh/AV-22-006245/2021 | 0.2532 | 0.0905 | 0.2232 | 0.0772 | 0.2 | Intermediate / Drift |
| 198 | A/broiler-chicken/Bangladesh/AV-22-079768/2021 | 0.2486 | 0.121 | 0.2054 | 0.0836 | 0 | Intermediate / Drift |
| 199 | A/broiler-chicken/Bangladesh/AV-22-006451/2021 | 0.2466 | 0.094 | 0.2177 | 0.0695 | 0 | Intermediate / Drift |
| 200 | A/sonali-chicken/Bangladesh/AV-22-079817/2021 | 0.2458 | 0.1059 | 0.2113 | 0.0984 | 0.2 | Intermediate / Drift |
| 201 | A/sonali-chicken/Bangladesh/AV-22-079757/2021 | 0.2446 | 0.0941 | 0.2165 | 0.012 | 0.3 | Intermediate / Drift |
| 202 | A/deshi-chicken/Bangladesh/AV-22-006643/2021 | 0.2384 | 0.0884 | 0.2158 | 0.0769 | 0 | Intermediate / Drift |
| 203 | A/deshi-chicken/Bangladesh/AV-22-079902/2021 | 0.2366 | 0.0905 | 0.2137 | 0.1408 | 0.1 | Intermediate / Drift |
| 204 | A/sonali-chicken/Bangladesh/AV-22-079790/2021 | 0.2361 | 0.0982 | 0.2096 | 0.0862 | 0 | Intermediate / Drift |
| 205 | A/sonali-chicken/Bangladesh/AV-22-006472/2021 | 0.2323 | 0.0992 | 0.2069 | 0.069 | 0.1 | Intermediate / Drift |
| 206 | A/sonali-chicken/Bangladesh/AV-22-006269/2021 | 0.2321 | 0.0695 | 0.2216 | 0.0099 | 0.1 | Intermediate / Drift |
| 207 | A/sonali-chicken/Bangladesh/AV-22-079795/2021 | 0.2253 | 0.0946 | 0.2052 | -0.1449 | 0 | Intermediate / Drift |
| 208 | A/sonali-chicken/Bangladesh/AV-22-079784/2021 | 0.2247 | 0.1002 | 0.2021 | 0.0682 | 0 | Intermediate / Drift |
| 209 | A/sonali-chicken/Bangladesh/AV-22-006265/2021 | 0.2135 | 0.0718 | 0.2098 | 0.0101 | 0.1 | Intermediate / Drift |
| 210 | A/broiler-chicken/Bangladesh/AV-22-079958/2021 | 0.1986 | 0.0802 | 0.1971 | -0.0656 | 0 | Intermediate / Drift |
| 211 | A/deshi-chicken/Bangladesh/AV-22-079956/2021 | 0.1922 | 0.0761 | 0.1955 | -0.1149 | 0.1 | Intermediate / Drift |
| 212 | A/deshi-chicken/Bangladesh/AV-22-006514/2021 | 0.1283 | 0.0717 | 0.1611 | 0.0925 | 0 | Intermediate / Drift |
| 213 | A/deshi-chicken/Bangladesh/AV-22-079866/2021 | 0.0973 | 0.0527 | 0.1528 | 0.0571 | 0.2 | Intermediate / Drift |
| 214 | A/deshi-chicken/Bangladesh/AV-22-079865/2021 | 0.0973 | 0.0527 | 0.1528 | 0.0571 | 0.3 | Intermediate / Drift |
| 215 | A/sonali-chicken/Bangladesh/AV-22-006636/2021 | 0.0898 | 0.1489 | 0.1007 | 0.103 | 0.1 | Intermediate / Drift |

**Table S3:**
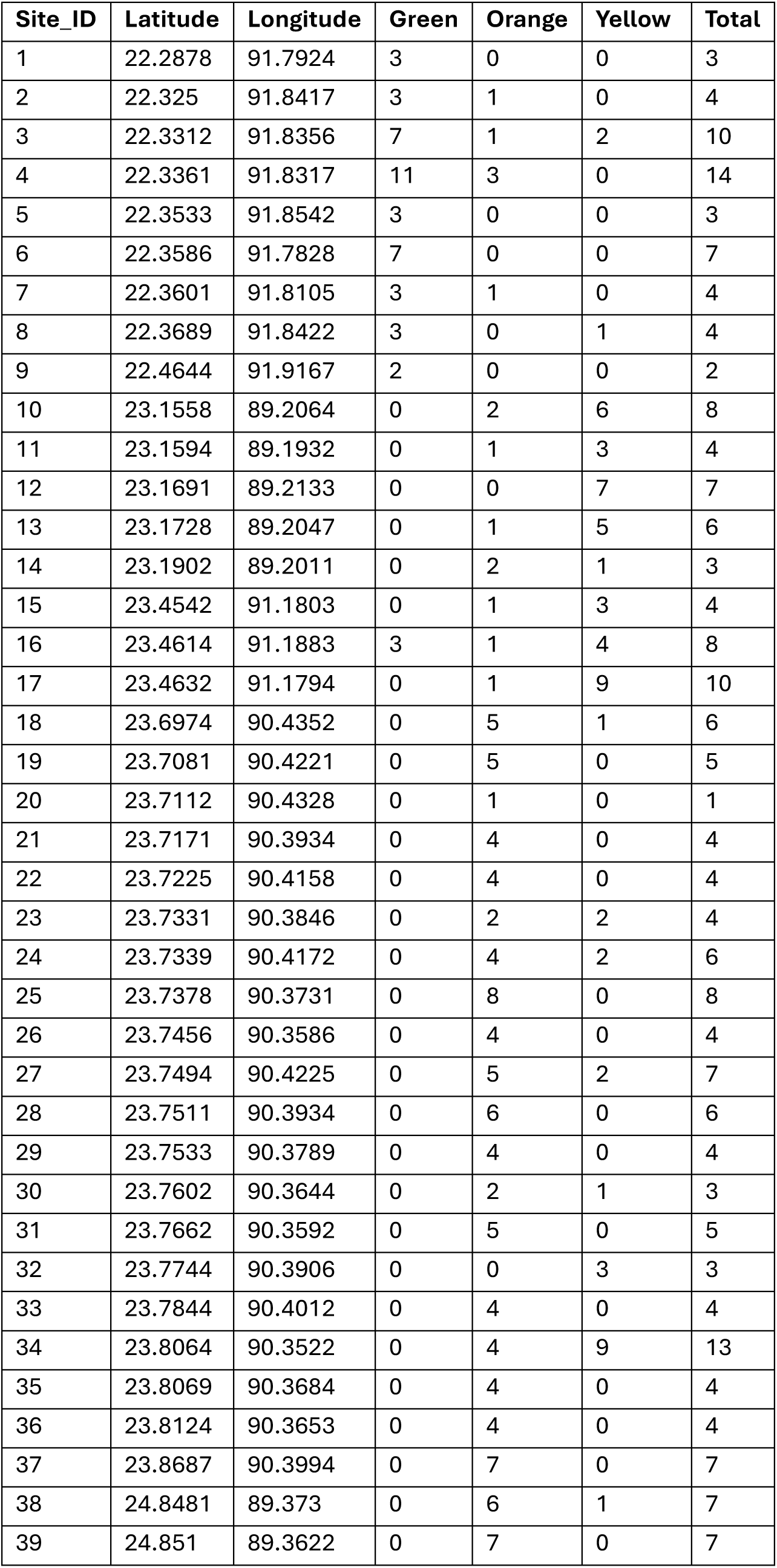
Bangladesh isolation site coordinates and frequency of isolation per site associated coloured by monophyletic group (Figure 1).

**Table S4:** N-linked glycan prediction of haemagglutinins (HA) and neuraminidases (NA) of sub-group representative strains using NetNGlyc 1.0 [64]. Glycan position (mature H9HA or H9N2 NA numbering), N-glycan motif (N-X-S/T), potential mean output score of nine neural networks, jury agreement of the nine neural networks and result (where +++ is >0.75, ++ is >0.6, + is >0.5 and - <0.5 confidence, thus deemed negative) values indicated per strain protein. Key substitutions between the sub-group representative strains are highlighted in bold.

| BNG924 (HA)<br>A/sonali-chicken/Bangladesh/AV-22-079924/2021 (yellow) |  |  |  |  |
|---|---|---|---|---|
| Position | Motif | Potential | Jury Agreement | Result |
| 11 | NSTE | 0.7339 | 9/9 | ++ |
| 64 | NPSC | 0.7547 | 9/9 | +++ (Proline) |
| 87 | NGTC | 0.8160 | 9/9 | +++ |
| 123 | NVTY | 0.7553 | 9/9 | +++ |
| 280 | NSTL | 0.7287 | 9/9 | ++ |
| 287 | **NISK** | 0.6997 | 9/9 | ++ |
| 474 | NGTY | 0.5836 | 7/9 | + |
| 533 | NGSC | 0.4602 | 3/9 | - |
| BNG943 (HA)<br>A/deshi-chicken/Bangladesh/AV-22-079943/2021 (green) |  |  |  |  |
| Position | Motif | Potential | Jury Agreement | Result |
| 11 | NSTE | 0.7338 | 9/9 | ++ |
| 64 | NPSC | 0.7408 | 9/9 | ++ (Proline) |
| 87 | NGTC | 0.8159 | 9/9 | +++ |
| 123 | NVTY | 0.7584 | 9/9 | +++ |
| 280 | NSTL | 0.7284 | 9/9 | ++ |
| 287 | **NVSK** | 0.7316 | 9/9 | ++ |
| 474 | NGTY | 0.5942 | 7/9 | + |
| 533 | NGSC | 0.4604 | 3/9 | - |
| BNG948 (HA)<br>A/sonali-chicken/Bangladesh/AV-22-079948/2021 (orange) |  |  |  |  |
| Position | Motif | Potential | Jury Agreement | Result |
| 11 | NSTE | 0.7338 | 9/9 | ++ |
| 64 | NPSC | 0.7308 | 9/9 | ++ (Proline) |
| 87 | NGTC | 0.8158 | 9/9 | +++ |
| 123 | NVTY | 0.7569 | 9/9 | +++ |
| 280 | NSTL | 0.7321 | 9/9 | ++ |
| 287 | **NISK** | 0.6997 | 9/9 | ++ |
| 474 | NGTY | 0.5941 | 7/9 | + |
| 533 | NGSC | 0.4604 | 3/9 | - |
| BNG924 (NA)<br>A/sonali-chicken/Bangladesh/AV-22-079924/2021 (yellow) |  |  |  |  |
| Position | Motif | Potential | Jury Agreement | Result |
| 29 | NSST | **0.5431** | **7/9** | **+** |
| 46 | NITE | 0.7494 | 9/9 | ++ |
| 54 | NGTI | 0.7649 | 9/9 | +++ |
| 71 | NWSK | 0.5706 | 8/9 | + |
| 131 | NGTT | 0.6942 | 9/9 | ++ |
| 185 | NATA | 0.3538 | 9/9 | -- |
| 219 | NGTC | 0.7598 | 9/9 | +++ |
| 387 | NRSG | 0.3548 | 9/9 | -- |
| BNG943 (NA)<br>A/deshi-chicken/Bangladesh/AV-22-079943/2021 (green) |  |  |  |  |
| Position | Motif | Potential | Jury Agreement | Result |
| 29 | NSST | 0.4762 | 4/9 | - |
| 46 | NITE | 0.7441 | 9/9 | ++ |
| 54 | **NGTV** | 0.7842 | 9/9 | +++ |
| 71 | NWSK | **0.6147** | **9/9** | **++** |
| 131 | **NDTT** | **0.5804** | **8/9** | + |
| 185 | NATA | 0.3550 | 9/9 | -- |
| 219 | NGTC | 0.7598 | 9/9 | +++ |
| 387 | NRSG | 0.3548 | 9/9 | -- |
| BNG948 (NA)<br>A/sonali-chicken/Bangladesh/AV-22-079948/2021 (orange) |  |  |  |  |
| Position | Motif | Potential | Jury Agreement | Result |
| 29 | NSST | 0.5423 | 7/9 | + |
| 46 | NITE | 0.7443 | 9/9 | ++ |
| 54 | NGTI | 0.7661 | 9/9 | +++ |
| 71 | NWSK | 0.5692 | 8/9 | + |
| 131 | NGTT | 0.6766 | 9/9 | ++ |
| 185 | NATA | 0.3536 | 9/9 | -- |
| 219 | NGTC | 0.7598 | 9/9 | +++ |
| 387 | NRSG | **0.4450** | **5/9** | **-** |

**Table S5:** Comparative TCID50, haemagglutinating units (HAU) and plaque-forming units (pfu)

| **Titres** | **Virus Name** |  |  |
|  | **BNG924** | **BNG943** | **BNG948** |
|---|---|---|---|
| **TCID50(/mL)** | 4.9x10<sup>6</sup> | 3.4x10<sup>7</sup> | 7.2x10<sup>6</sup> |
| **HAU (/50µL)** | 512 | 256 | 512 |
| **Plaque (pfu/mL)** | 2.0x10<sup>8</sup> | 1.2x10<sup>8</sup> | 8.1x10<sup>7</sup> |
| **Plaque Morphology** |  |  |  |

**Figure S1:**
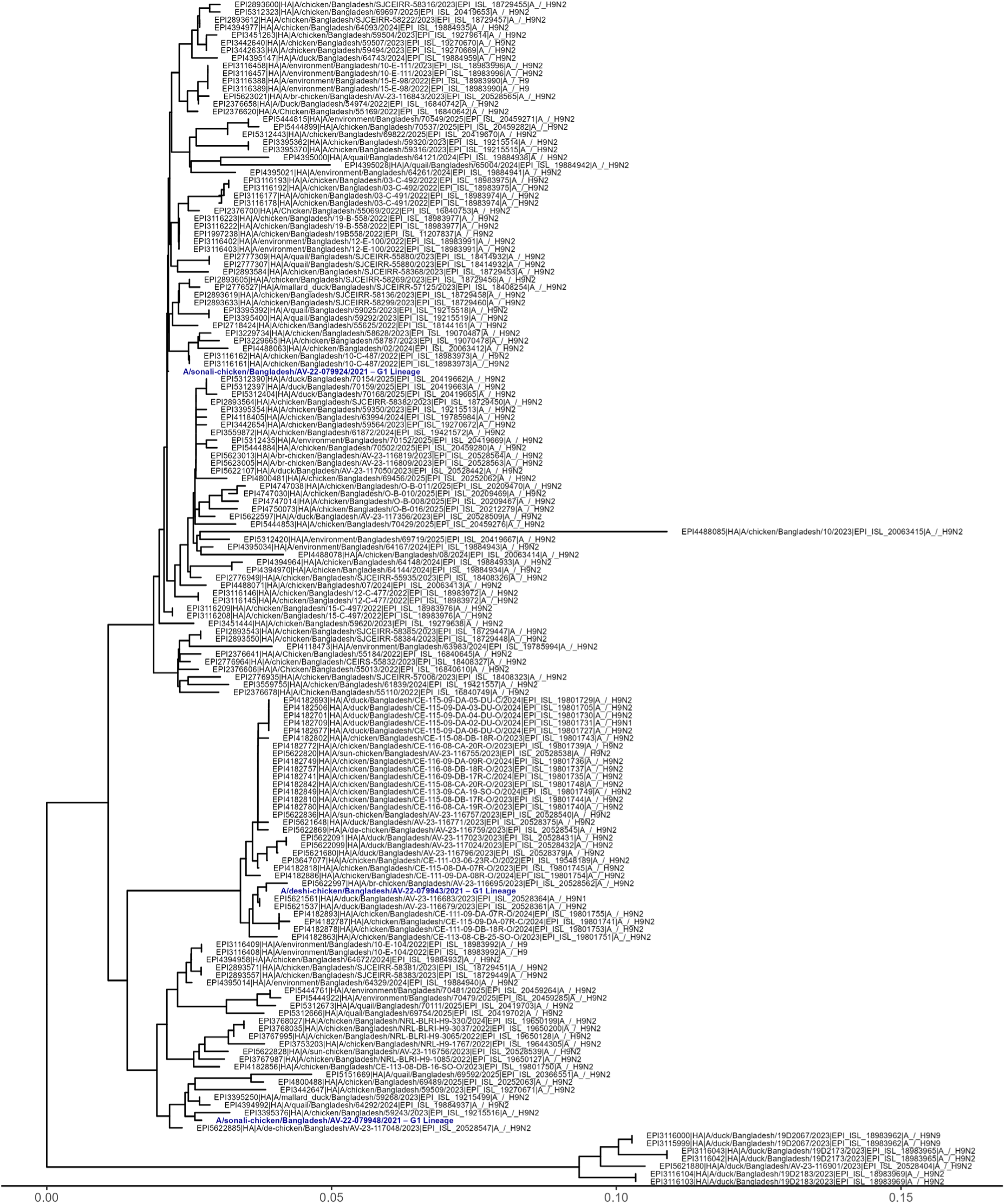
Neighbour-Joining phylogenetic tree of trimmed mature haemagglutinin (HA) amino acid sequences (n = 155) of isolates collected from Bangladesh between 2022-2026 from GISAID database. Scale indicates rate of genetic distance with 0.01 corresponding to 1% difference in amino acid sequence. Tree generated in RStudio [15] with ape [16], ggtree [18], ggplot2 [65] & tidyverse [19] packages.

